# Microbial cancer immunotherapy reprograms hematopoiesis to enhance anti-tumor immunity

**DOI:** 10.1101/2024.03.21.586166

**Authors:** Andrew W. Daman, Anthony Claude Antonelli, Gil Redelman-Sidi, Lucinda Paddock, Jin Gyu Cheong, Leonardo F. Jurado, Anna Benjamin, Song Jiang, Dughan Ahimovic, Shireen Khayat, Michael J. Bale, Oleg Loutochin, Victor A. McPherson, Dana Pe’er, Maziar Divangahi, Eugene Pietzak, Steven Z. Josefowicz, Michael S. Glickman

## Abstract

*Mycobacterium bovis* BCG is the vaccine against tuberculosis and an immunotherapy for bladder cancer. When administered intravenously, BCG reprograms bone marrow hematopoietic stem and progenitor cells (HSPCs), leading to heterologous protection against infections. Whether HSPC-reprogramming contributes to the anti-tumor effects of BCG administered into the bladder is unknown. We demonstrate that BCG administered in the bladder in both mice and humans reprograms HSPCs to amplify myelopoiesis and functionally enhance myeloid cell antigen presentation pathways. Reconstitution of naive mice with HSPCs from bladder BCG-treated mice enhances anti-tumor immunity and tumor control, increases intratumor dendritic cell infiltration, reprograms pro-tumorigenic neutrophils, and synergizes with checkpoint blockade. We conclude that bladder BCG acts systemically, reprogramming HSPC-encoded innate immunity, highlighting the broad potential of modulating HSPC phenotypes to improve tumor immunity.

**Graphical Abstract:** 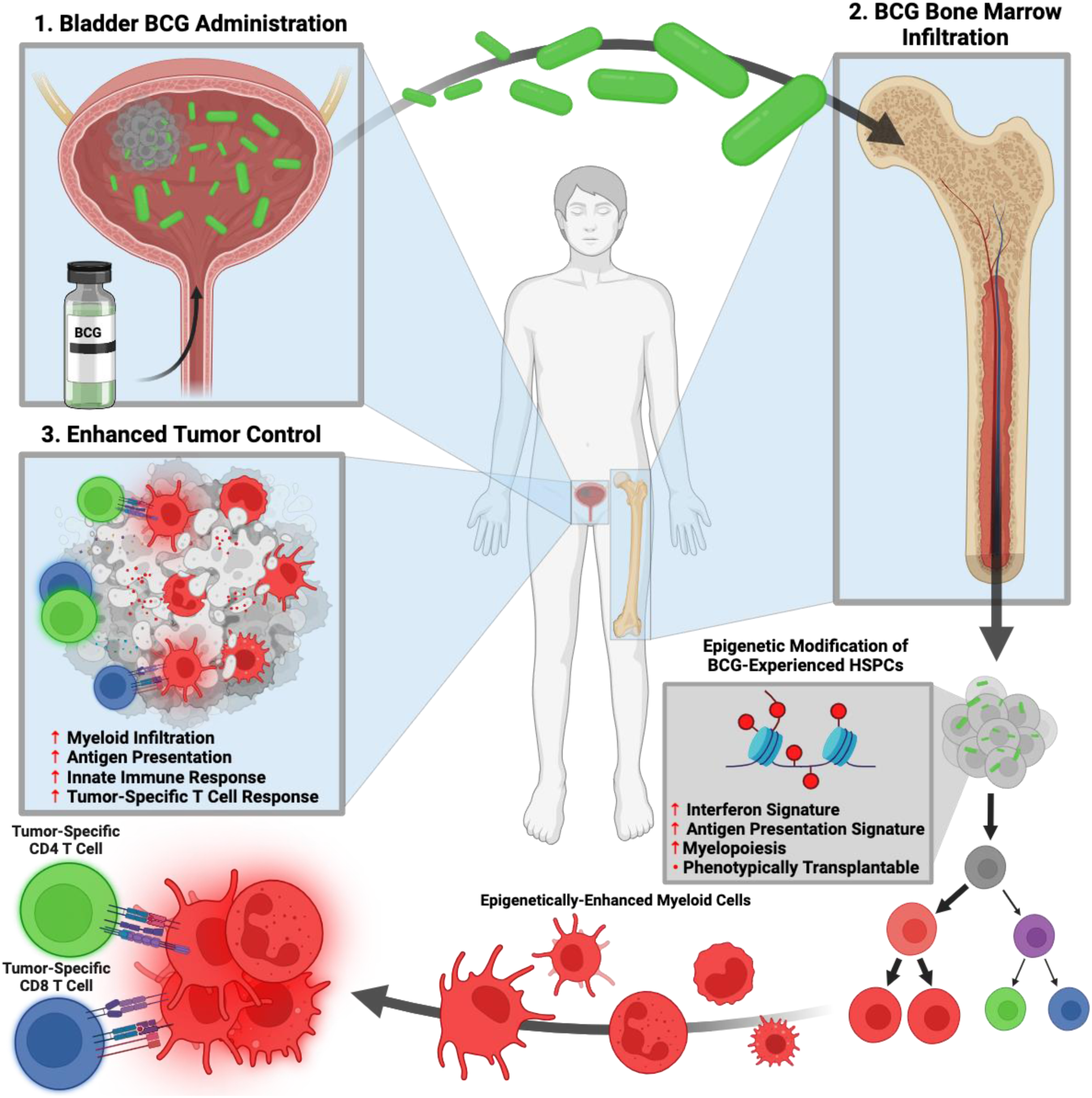

## Introduction

Bacillus Calmette-Guérin (BCG) is an attenuated strain of *Mycobacterium bovis* used worldwide as a live vaccine for tuberculosis. BCG is also the first cancer immunotherapy (1) and the only bacterial therapy for cancer. Intravesical BCG therapy, the instillation of live BCG directly into the bladder, is the standard of care for non-muscle-invasive bladder cancers (NMIBC) (2) due to its ability to reduce recurrence rates and improve overall survival when compared with surgical resection alone (3–6). However, approximately 50% of bladder cancer patients fail to maintain durable responses to BCG therapy (7–10), and there are no well-established biomarkers to predict responses in advance of treatment initiation. This is in part due to an incomplete understanding of the mechanism by which BCG mediates tumor clearance (11,12).

Following bladder instillation, BCG attaches to urothelial cells (13–16), resulting in recruitment and tumor infiltration of myeloid and lymphoid cells in both humans and mice (17,18). In mouse models, bladder tumor rejection is mediated by tumor specific CD4 and CD8 T cell immunity (19–22). Complementary evidence from human studies has identified tumor-specific CD4 T cells in BCG-treated patients with NMIBC (23). These effects of BCG are presumed to be mediated locally within the bladder, but the upstream events stimulated by BCG that enable tumor-specific immunity remain poorly defined.

An expanding body of research has established that microbes and their products, including BCG, can lead to epigenetic changes in innate immune cells that confer an increased capacity to respond not only to homologous but to heterologous immune stimuli, a process termed innate immune memory or trained immunity (24–26). The persistence of these phenotypes in short-lived innate immune cells is explained by microbe-induced epigenetic changes in hematopoietic stem and progenitor cells (HSPCs) in the bone marrow (24,27). Epigenetic changes in HSPCs (central innate immune memory) are associated with skewed hematopoiesis toward myeloid cell production (myelopoiesis) and can be passed to progeny cells, poising mature myeloid cells for heterologous immunity (28, 29). Single-cell analysis of gene expression programs in human monocytes following BCG vaccination revealed that increased activity of type II *versus* type I interferon programs are linked to augmented innate immune function (29,30). This type of induced and durably augmented innate immune activity can confer heterologous protection against infection, including against respiratory viruses in BCG vaccinated children or elderly nursing home residents (31–34). Both the fungal cell wall component beta-glucan(35) and a synthetic bone marrow targeted peptidoglycan (36) have been shown to augment anti-tumor immunity in mouse models of melanoma *via* myeloid-reprogramming, highlighting the potential of tuning these pathways for augmenting anti-tumor responses. Although intravenous BCG administration also generates innate immune memory at the level of HSPCs (27), it is unknown whether HSPC-reprogramming occurs upon BCG administration in the bladder and contributes to anti-tumor immunity. Recent studies indicate that bladder administration of BCG can increase circulating cytokines (37), modify monocyte phenotypes, and reduce frequency of upper respiratory infection (38). The relation of these observations to anti-tumor immunity is unknown, as are the mechanisms involved, including HSPC alterations. We sought to understand how bladder BCG, as a long-standing human immunotherapy, contributes to anti-tumor immunity, including if it acts systemically to reprogram HSPCs and if so, how this reprogramming contributes to a functional anti-tumor response. We demonstrate that bladder BCG reprograms HSPC, which pass on augmented antigen presentation and migratory potential to mature innate immune cells, and increase anti-tumor immune responses. These studies highlight the potential of tuning HSPC phenotypes for anti-tumor immunity.

## Results

### Bladder BCG induces central innate immune memory

Intradermal administration of BCG in an adult human cohort (29,39) and intravenous administration in mice (27) have been shown to induce innate immune memory programs in HSPCs with prominent interferon-gamma (IFN-γ) signatures. In contrast, administration of BCG into the bladder has long been presumed to act locally by modifying the tumor microenvironment (12,40–42). To determine if bladder BCG stimulates innate immune memory in HSPCs of human NMIBC patients, we collected blood samples from two independent longitudinal post-resection cohorts (Memorial Sloan Kettering Cancer Center, n=8; and McGill University, n=13) before initial treatment and one week after the 5th administration of BCG instillation into the bladder (Figure 1A, Supplementary Table 1, Methods).

**Figure 1:**
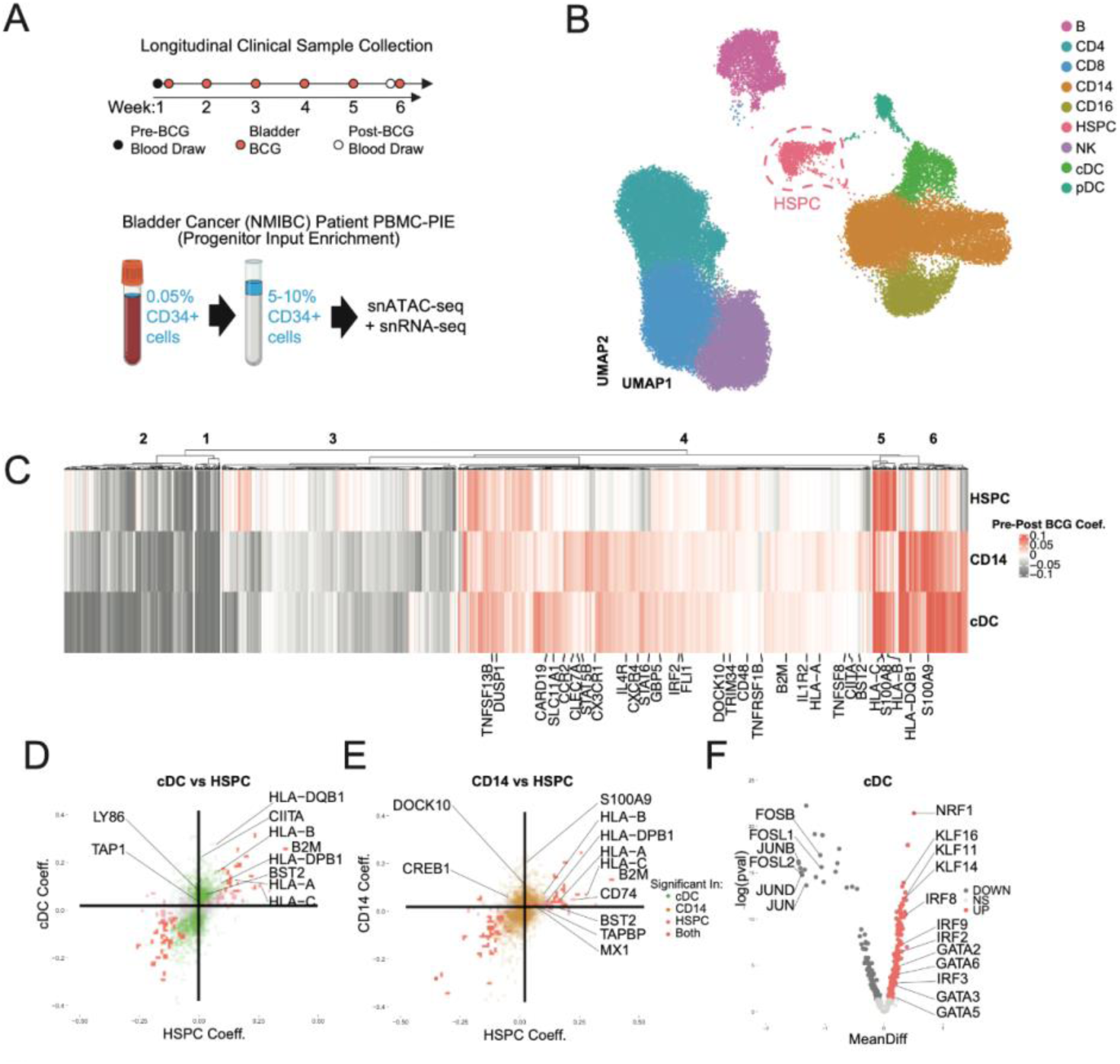
Bladder BCG reprograms HSPCs and myeloid progeny in human bladder cancer patients. A) Experimental Schematic. Whole blood was collected from non-muscle-invasive bladder cancer patients prior to their first dose of BCG and after their 6th dose of BCG (top). Cryopreserved PBMCs were analyzed by PBMC-PIE (40), which enriches rare HSPCs from peripheral blood, followed by mixing with the PBMC at a greater ratio (bottom). Paired single nucleus ATAC- and RNA-seq were performed on all samples. B) UMAP visualization of snRNA data from the experiment described in A, colored by cell type annotation, with the PBMC-PIE enriched HSPC population highlighted C) Fold change in expression post-*versus* pre-BCG for all significant differentially expressed (fdr < 0.01) genes in HSPCs, CD14+ monocytes, or conventional dendritic cells (cDCs). Coef. references the coefficient of change in expression in the MAST differential expression model. Heatmap is clustered using k-means and immunologically relevant genes are labeled. D) Scatterplots of fold change in expression for all genes post-*versus* pre-BCG in HSPCs *versus* conventional dendritic cells. Points are colored by significance (fdr <0.01) in one cell type or both, and relevant genes are labeled. E) Scatterplots of fold change in expression for all genes post-*versus* pre-BCG in HSPCs *versus* CD14+ monocytes. Points are colored by significance (fdr <0.01) in one cell type or both, and relevant genes are labeled. F) Volcano plot of differential chromVAR motif accessibility. MeanDiff is the difference in average chromVAR score for accessible peaks in conventional dendritic cells post- *versus* pre- BCG, and p-values were determined by Wilcoxon rank-sum test. Significant motifs (p < 0.05) are colored by direction of change.

We interrogated the bladder BCG-induced changes to HSPCs and mature immune cells by employing Peripheral Blood Mononuclear Cell analysis with Progenitor Input Enrichment (PBMC-PIE), a workflow recently established by our group (Figure 1A). This workflow allows for combined single nuclei RNA and ATAC sequencing of human HSPCs *via* isolation and enrichment of rare circulating CD34+ cells in the blood, which we have shown faithfully captures the extensive diversity and phenotypes of bone marrow CD34+ cells (43). Using PBMC-PIE and MEIS1 as a well-defined HSPC marker gene alongside a panel of standard immune cell type markers (44), we captured a total of 58,652 cells (57,543 from mature peripheral blood immune cell types and 1,118 circulating HSPCs, (Figure 1B, Figure S1A).

We first examined our two independent cohorts separately. In both cohorts, we found post-BCG transcriptional upregulation of genes associated with antigen presentation and interferon signaling in both HSPCs and mature myeloid cell progeny (Figure S1B-C). Given the degree of overlap in transcriptional changes between the two datasets, we applied an entropy-based measure to evaluate batch effect in combining cohorts (see methods) and found samples to be well mixed in RNA local cell neighborhoods (Figure S1D-E). Because our subsequent analysis is based on cell type clusters derived from the RNA neighbor graph, we did not apply further batch correction methods.

To identify molecular programs associated with central innate immune memory, we focused on analysis of the HSPC (MEIS1+) cluster and mature myeloid populations, namely CD14+ monocytes (CD14+, LYZ+) and classical dendritic cells (cDC) (FCER1A+, CST3+). Analysis of all significantly differentially expressed genes pre- and post-BCG therapy in either HSPC, CD14+ monocyte, or cDC compartments revealed prominent and broadly consistent changes in transcription following bladder BCG treatment (Figure 1C). In HSPCs, we observed significant post-BCG transcriptional upregulation of genes and pathways associated with antigen presentation (HLA-C, HLA-DRB5, HLA-DRA, HLA-DQB1, CD74 (invariant chain), and B2M), as well as genes with other immune-related functions such as ZEB2, an ETS family member and key regulator of myeloid development (35,45,46), and BST2, a regulator of HSPC activation downstream of IFN-γ (47) (Figure 1C-E). Of particular interest, when these significantly differentially expressed genes were clustered according to fold-change in expression, many of the genes associated with antigen presentation co-clustered in Clusters 4 and 5 (Figure 1C), and are shared across the three cell types, suggesting a shared antigen presentation program that persists through differentiation from HSPC to monocyte and cDC progeny. To visualize this shared program, we compared the fold change of significantly differentially expressed genes pre- and post-BCG in HSPCs with the corresponding fold change in either cDCs or monocytes. This visualization highlights key genes associated with interferon-gamma-mediated signaling and antigen presentation (Figure 1D-E), a finding that was confirmed by GO pathway analysis, with top enriched categories “interferon gamma mediated signaling pathway” (GO:0060333, p-value: 0.004) and “antigen processing and presentation of exogenous peptide antigen *via* MHC-I, TAP-independent” (GO:002480, p-value: 0.002) (Fig S1E). Overall, these data indicate that bladder BCG induces reprogramming of HSPCs and mature myeloid cells in humans, with changes characteristic of interferon-dependent pathways. Furthermore, the extensive transcriptional changes in HSPCs, remote from the mucosal site of administration, indicates that the effects of bladder BCG are systemic and not limited to the tumor microenvironment.

We next explored predicted transcription factor (TF) activity in HSPCs and mature immune cells that may be driving the altered transcription programs observed above, including those associated with augmented expression of antigen-presentation and IFN-γ response programs. In cDCs, we observed significant enrichment of characteristic interferon response factor (IRF) family motif accessibility post-BCG, as well as GATA and KLF motifs which have previously been associated with cellular survival and activation (48–50)(Figure 1F, S1F,G). In HSPCs, we observed significant enrichment for the predicted activity of AP-1 (FOS/JUN), RUNX (51,52), and TAL/ZEB/ETS family members (53–55) (Figure S1G). AP-1 has previously been shown to be associated with the formation of stem cell innate immune memory (43,43,56), and the strong association of ETS family members with myelopoiesis indicates that these HSPCs are reprogrammed for increased myeloid output post-BCG.

Collectively, these findings establish a BCG exposure signature in both HSPCs and their mature myeloid cell progeny (monocyte and cDC) following bladder administration of BCG in humans; both chromatin accessibility and transcriptomic analyses indicate a signature consistent with interferon imprinting. The data further indicates that the previously characterized systemic effect of intravenous BCG on HSPC-driven innate immune memory is also a feature of BCG administered in the bladder as an immunotherapy for cancer.

### Bladder BCG alters HSPC composition *via* direct bone marrow colonization

We next sought to address whether BCG induced innate immune memory directly contributes to anti-tumor immunity. We employed the MB49 murine model of bladder cancer, in which the syngeneic bladder tumor cell line MB49 is implanted into the bladder, followed by 5 weekly bladder installations of 3x10^6^ CFU of live BCG. In this model, 20% to 50% of BCG-treated mice demonstrate long-term survival and subsequent tumor immunity, while 100% of control (PBS-treated) mice succumb to disease. We found that intravenous BCG, known to provoke central innate immune memory in mice, provided similar levels of protection to bladder BCG, and there was no synergy when combining the two routes of administration (Figure 2A, 2B, Supplemental Figure 2B). To test whether bladder BCG similarly disseminates to bone marrow, as has been reported with intravenous BCG (27), we harvested and cultured bone marrow at weekly intervals during a 5-week course of bladder BCG administration. We found live BCG from all mice that had received the full 5 doses of bladder BCG, and several of the mice that had received 3 or 4 doses (Figures 2C, S2B).

**Figure 2:**
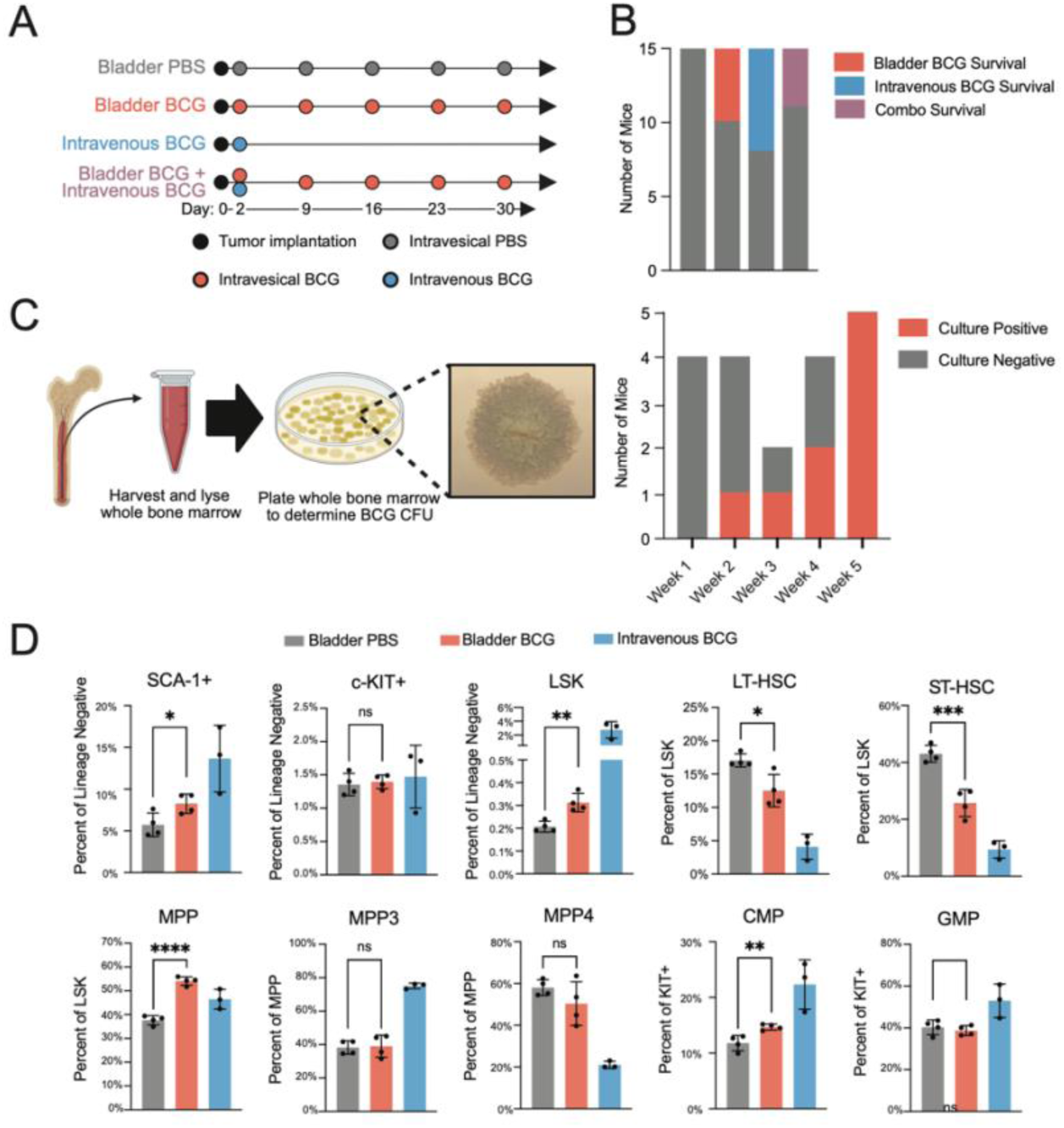
Bladder BCG reprograms bone marrow HSPCs *via* direct colonization. A) Experimental schematic. Mice were implanted with MB49 bladder tumors on Day 0 and administered one of the following regimens beginning on Day 2: 5 weekly doses of bladder PBS, 5 weekly doses of bladder BCG, a single dose of intravenous BCG, or a combination of single-dose intravenous BCG and 5 weekly doses of bladder BCG. Survival was monitored throughout. B) Quantification of overall survival from the experiment described in A. C) Bone marrow was harvested from mice treated with 1, 2, 3, 4, or 5 weekly doses of bladder BCG and cultured on agar medium to quantify live BCG colonization, as depicted on the left. The entirety of the femur and tibias from each mouse were harvested and lysed then plated for BCG growth. The proportion of culture positive *versus* culture negative samples is displayed according to weeks of BCG treatment on the right. Quantification of the number of colonies in each mouse bone marrow is provided in Supplemental Figure 2. D) Mice were administered 5 doses of bladder PBS, 5 doses of bladder BCG, or a single dose of intravenous BCG. Bone marrow was harvested 24 hours after the third dose of bladder PBS or bladder BCG and HSPC subsets were quantified by flow cytometry. Intravenous BCG samples were from an independent reference experiment and were not compared statistically with bladder PBS or bladder BCG samples. P values were derived by Student’s t-test. P > 0.05 = ns, P ≤ 0.05 = *, P ≤ 0.01 = **, P ≤ 0.001 = ***, P ≤ 0.0001 = ****.

Prior studies have established that intravenous BCG induces expansion and epigenetic modification of the lineage^-^ Sca-1^+^ c-Kit^+^ (LSK) population of HSPC linked to the functional enhancement of mature myeloid cells (27,29,57). Flow cytometric analysis of HSPC subsets (Fig S2C) after bladder BCG revealed expansion of the LSK population, decreases in the proportion of long-term hematopoietic stem cells (LT-HSCs) and short-term hematopoietic stem cells (ST-HSCs), an increase in the proportion of multipotent progenitor (MPP) cells, and an increase in common myeloid progenitor cells (CMPs) (Figure 2D). To further define the importance of the route of administration to this phenomenon, we also profiled the effect of subcutaneous BCG, and in line with previously published results (27), subcutaneous BCG showed a reduced increase in LSK frequency compared with bladder and intravenous administration (Figure S2D). Taken together, these results indicate that BCG administered in the bladder colonizes the bone marrow and alters hematopoiesis by increasing immune cell production skewed toward the myeloid lineage, a hallmark of innate immune memory.

### Bladder BCG remodels the HSPC chromatin landscape

To determine the cellular and molecular programs that underlie HSPC expansion and inflammatory hematopoiesis following bladder BCG, we performed in-depth single-cell epigenomic and transcriptomic analyses of HSPCs from mice bearing MB49 bladder tumors after 3 doses of bladder BCG (Figure 3A). After identifying HSPC and mature cell subsets based on marker genes (Figure 3B, Figure S3A), we compared the distribution of cells from BCG- and PBS-treated mice and observed a notable increase in the density of cells in the neutrophil progenitor cluster in BCG-treated mice (Figure 3C), indicating that a component of the myeloid skewing induced by bladder BCG includes the neutrophil lineage. Next, to define epigenomic-reprogramming in HSPCs, we focused further analyses on the snATAC-seq data (we avoided integration and in-depth analyses of snRNA because we observed low read depth in our snRNA libraries, a common feature of bone marrow multiome; communications with 10X Genomics). Predicted TF activity analysis (motif accessibility by ChromVAR) revealed enriched IRF and STAT TF activity across stem cells and myeloid progenitors (HSC/MPP, neutrophil progenitor, and monocyte progenitor) (Figure 3D), strongly indicating that bladder BCG causes a systemic response that exposes HSPCs to interferon signaling, similar to what has been characterized with intravenous administration of BCG (27). The presence of these changes in more differentiated progenitor populations led us to hypothesize the persistence of epigenetic features through differentiation with functional consequences in fully mature myeloid cells (see below). Differential gene expression analysis supported these observations with transcriptional changes in HSC:MPP, neutrophil progenitor, and monocyte progenitor populations consistent with an interferon-responsive antigen presentation signature (Figure S3B).

**Figure 3:**
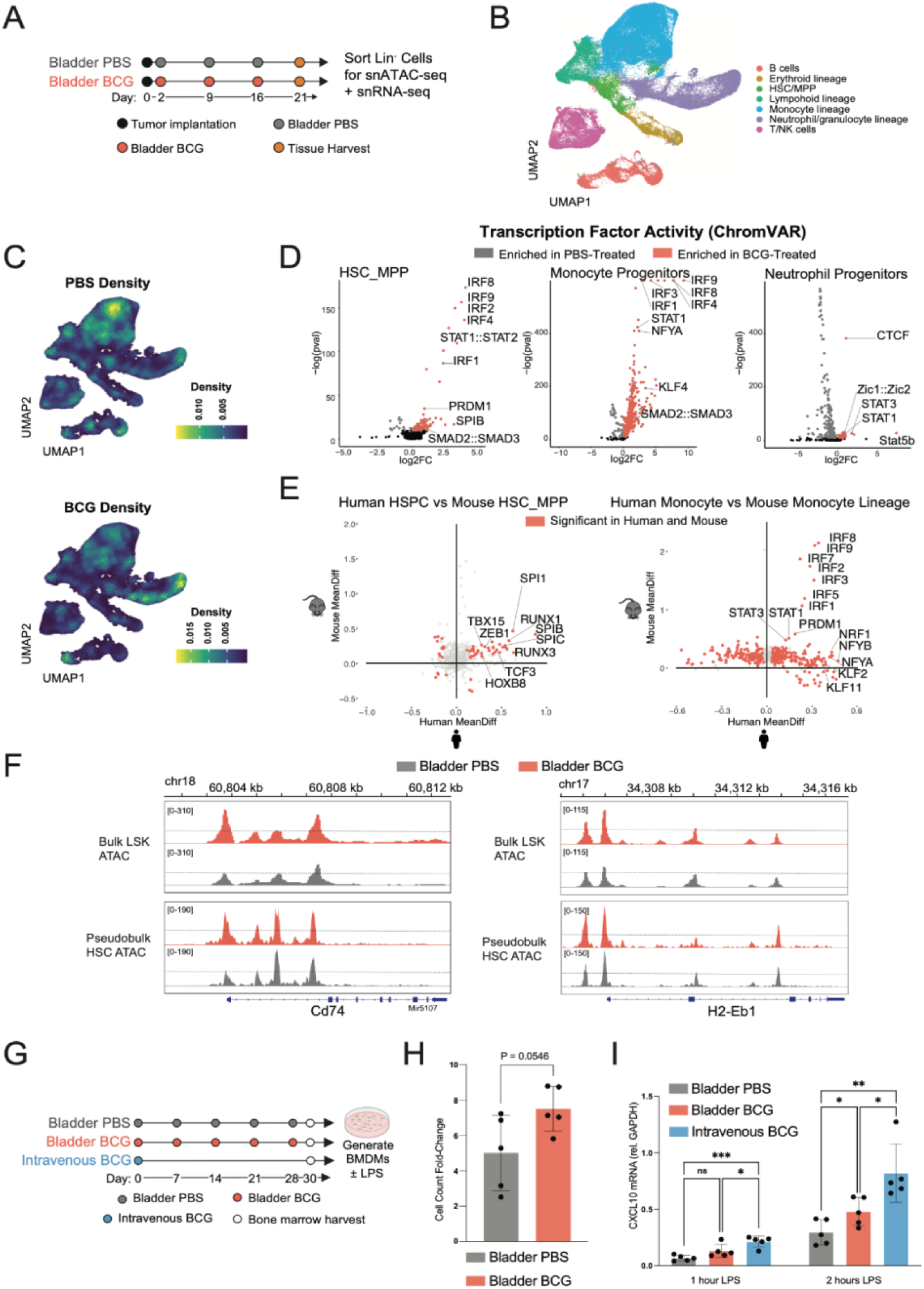
BCG modulates the differentiation potential of bone marrow HSPCs and enhances the function of their myeloid progeny. A) Experimental schematic. Mice were implanted with MB49 tumors on Day 0 and administered 3 weekly doses of bladder PBS or bladder BCG beginning on Day 2. Bone marrow was harvested 7 days after the final administration, sorted for Lineage-cells (CD3-, B220-, Ter119-, GR1-, NK1.1-), and paired single nucleus ATAC- and RNA-seq was performed. B) UMAP showing diverse cellular lineages from the experiment described in A. Cells were annotated using cell type references from SingleR. C) Density of PBS- or BCG-treated cells projected onto the UMAP, showing differential density in the monocyte and neutrophil precursor lineages in BCG-treated mice. D) Volcano plots depicting predicted differential transcription factor activity inferred from snATAC-seq for HSC/MPP, monocyte, and neutrophil populations. Red indicates significant enrichment in BCG-treated cells while gray indicates significant enrichment in PBS-treated cells. E) Correlation plots depicting conserved predicted differential transcription factor activity between human and mouse cell subsets. Human HSPC *versus* mouse HSC/MPP populations are shown on the left, human monocytes *versus* mouse monocytes are shown on the right. Regions of significant predicted transcription factor activity common to both humans and mice are highlighted in red. F) Comparison of ATAC-seq tracks generated from bulk ATAC-seq of sorted LSK cells (Lineage-Sca-1+ c-Kit+ cells) and pseudo-bulk ATAC-seq generated by concatenating all ATAC-seq reads from the HSC/MPP cluster in the single nucleus bone marrow data. Comparisons for two significant differentially accessible chromatin regions associated with antigen presentation are displayed: CD74 (left) and H2-Eb1 (right). G) Experimental schematic. Mice were given 5 weekly doses of bladder PBS, bladder BCG, or a single dose of intravenous BCG. Bone marrow was harvested 1 week after the final bladder treatment and bone marrow-derived macrophages (BMDMs) were generated. BMDMs were stimulated with LPS after 10 days of differentiation. H) Yield of BMDMs. Total cells were counted at Day 0 (prior to BMDM differentiation) and Day 10 (after terminal differentiation into BMDMs). Cell count fold change is plotted for BMDMs from bladder PBS- and bladder BCG-treated mice. I) BMDMs were stimulated with LPS for 1 or 2 hours. RNA was extracted and qPCR was performed for the chemokine CXCL10. Data is plotted relative to GAPDH. P values were calculated by Student’s t-test. P > 0.05 = ns, P ≤ 0.05 = *, P ≤ 0.01 = **, P ≤ 0.001 = ***, P ≤ 0.0001 = ****.

To determine the concordance of the mouse and human BCG-induced reprogramming, we co-visualized myeloid cell TF activity programs from both our mouse and human datasets, plotting the fold change of each orthologous chromVAR score and denoting those that were significantly different in both species (Figure 3E). In HSC/MPP we saw a consistent activation of TFs that regulate myelopoiesis (PU.1, RUNX1, Zeb1), and in monocytes and dendritic cells, we saw a consistent activation of IRF family members and STAT1/3 (Figure 3E). We also observed concordant mouse and human RNA upregulation of antigen presentation and IFN-γ responses in both monocytes and dendritic cells (Figure S3C).

To confirm these results in a complementary model and with a purified bulk stem cell population, we treated mice with 5 doses of either bladder PBS or BCG and performed bulk ATAC sequencing on sorted LSK cells (Figure S3D). Principal Component Analysis revealed clustering of LSKs from BCG-treated replicates compared to PBS-treated mice, with PC1 capturing chromatin accessibility associated with BCG treatment (Figure S3E). Differential peak accessibility analysis showed an overall upregulation of accessibility after BCG treatment (Figure S3F). Consistent with our findings from human and mouse single cell analysis, HOMER motif analysis of differential peak accessibility from bulk ATAC sequencing revealed increased inferred TF activity for IRF family members, along with NFY, a TF required for MHC enhanceosome formation and transcription of MHC-II genes (Figure S3F) (58), and PU.1 (Spi1), a master regulator of hematopoiesis crucially important for myeloid cell development (59). Comparison of pseudobulk tracks from the HSC/MPP snATAC-seq data with LSK ATAC-seq (Figure 3F) revealed concordance across single cell annotations of HSC/MPP with sorted LSK, and between these different experiments, with extensive similarities, including at genes involved in antigen presentation, such as CD74, and H2-Eb1 (Fig 3F).

### Bladder BCG activates HSPC-encoded macrophage hyper-responsiveness

We next asked if the population-level and epigenetic changes stimulated by BCG in HSPCs and their myeloid progeny result in functional enhancement by comparing bone marrow-derived macrophages (BMDMs) from mice treated with bladder BCG *versus* PBS. To establish that any functional changes originate from epigenetic programs in early stem cells, we FACS-sorted LSK cells from bone marrow before inducing macrophage differentiation *in vitro* (Figure 3G). Consistent with the increased myelopoiesis documented above in BCG-treated mice, we saw a trend toward greater *in vitro* expansion of LSK-derived macrophages from mice treated with bladder BCG (Figure 3H). Stimulation of LSK-derived macrophages with LPS revealed significant enhancement of transcripts encoding the T cell-recruiting chemokine CXCL10 in macrophages derived from both bladder- and intravenous BCG-treated LSKs compared to control macrophages (Figure 3I, S3H).

### BCG-experienced HSPCs are sufficient to limit tumor growth

We next sought to determine the contribution of BCG induced epigenetic alterations in HSPCs to mature myeloid phenotypes and tumor control. We transplanted bone marrow from bladder or intravenous BCG-treated CD45.2^+/+^ donor mice into naive irradiated CD45.1^+/+^ recipient mice (Figure 4A). Flow cytometric analysis of PBMC subsets from bone marrow chimeric mice at 8-weeks post-transplant confirmed complete immune reconstitution, with approximately 95% of circulating immune cells of donor origin. In line with our previous observations of post-BCG myeloid skewing, and demonstrating the HSPC origin of these changes, we observed an increase in circulating myeloid cells derived from BCG-experienced donor bone marrow *versus* PBS controls (Figure 4B). To assess the epigenetic programs and phenotypes of defined progenitors more directly, we designed parallel experiments with chimeric animal protocols with sorted LSK populations rather than total bone marrow. While settings of inflammation can alter LSK cell surface marker expression (including increasing Sca-1) (60), we mitigated bias and heterogeneity of sorted LSK populations by culturing them in selective, primitive HSC-expanding media conditions (61) for three weeks before transplant.

**Figure 4:**
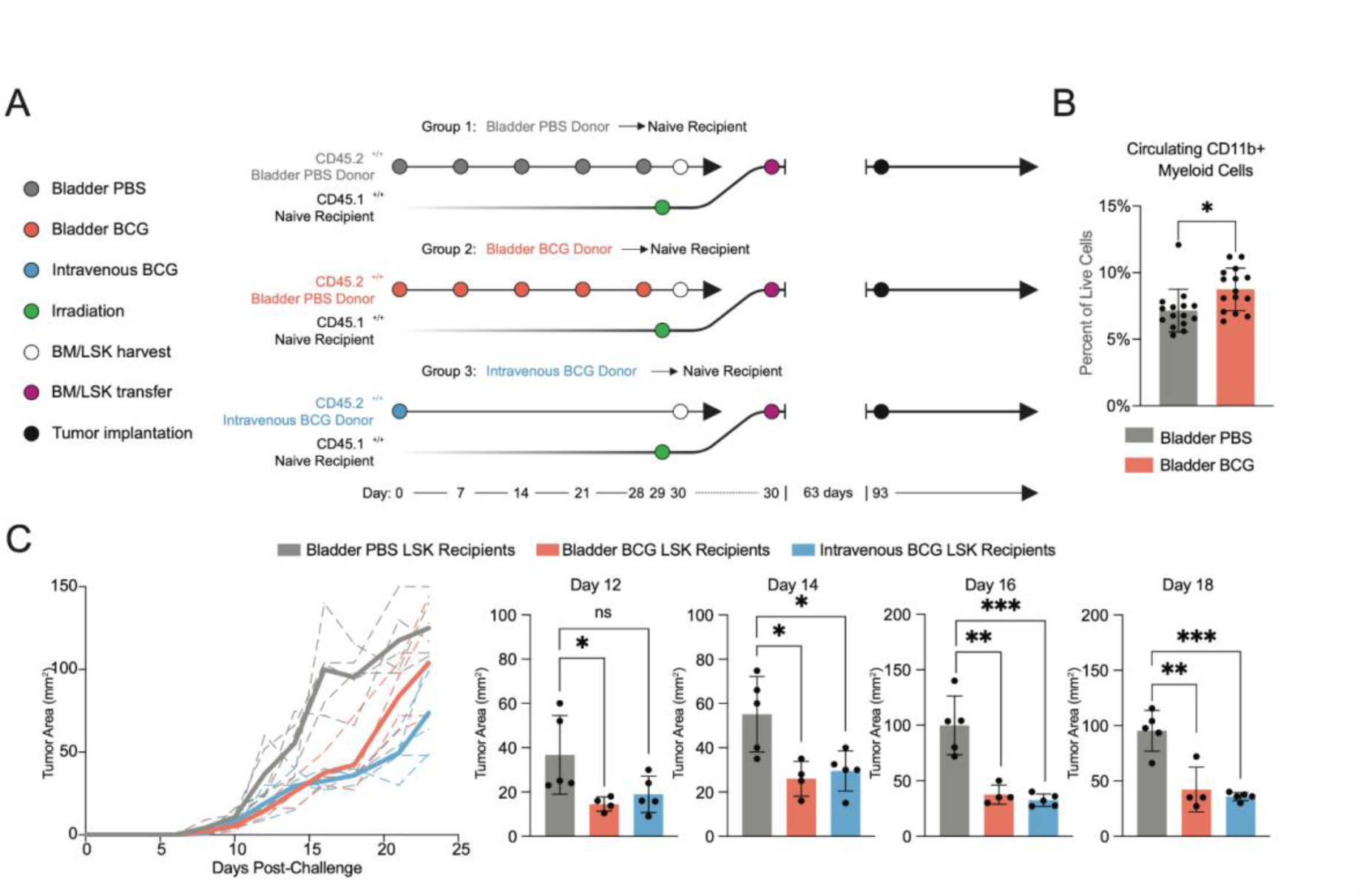
BCG-experienced HSPCs confer enhanced control of tumors to naive recipient mice. A) Experimental schematic. Congenically-marked bone marrow chimeras were generated by transferring sorted bone marrow LSK cells from CD45.2^+/+^ donor mice treated with 5 weekly doses of bladder PBS, 5 weekly doses of bladder BCG, or single-dose intravenous BCG, into CD45.1^+/+^ naive irradiated recipient mice. After confirmation of full immune cell reconstitution from donor LSK cells, mice were challenged with subcutaneous MB49 tumors and growth was monitored longitudinally. B) Enhancement of bone marrow myeloid output by BCG-reprogrammed HSPCs. CD45.1^+/+^ naive irradiated mice were transferred bulk bone marrow from bladder PBS- or bladder BCG-treated CD45.2^+/+^ donor mice. After 8 weeks, circulating myeloid cells were quantified. C) Tumor growth curves from the experiment described in A (left). Dotted lines represent individual mice within a group, solid lines represent the group average. Statistical comparisons for Days 12, 14, 16, and 18 are displayed (right). P values were derived by Student’s t-test. P > 0.05 = ns, P ≤ 0.05 = *, P ≤ 0.01 = **, P ≤ 0.001 = ***, P ≤ 0.0001 = ****.

This system has the additional benefits of allowing for inflammatory programs to resolve and reducing potential for transfer of live BCG along with LSK cells which could induce training in the recipient mouse (Intravenous administration of BCG does not result in direct infection of LSK cells, and rather, the BCG resides in CD11b+ Ly6C+ monocytes (27)).

Chimeric animals generated both from cultured LSK (Figure 4C) and from whole bone marrow (Figure S4A) were then implanted with subcutaneous tumors. BCG exposed donor LSKs conferred enhanced control of tumor growth, with no discernible difference between the two routes of BCG administration. Challenge of BCG LSK-reconstituted mice with B16 melanoma tumors revealed a similar effect on tumor control (Figure S4B) establishing that the tumor control conferred by BCG-reprogrammed HSPCs is tumor-type and presumably antigen-independent. As an additional control to confirm that the tumor control conferred by transplanted LSKs is not the result of transfer of viable BCG, we treated donor LSK cells with the anti-mycobacterial antibiotic isoniazid prior to transfer into recipient mice and observed similar tumor control (Figure S4C). These results confirm that the LSK HSPC subset alone, without contribution from downstream progenitors or other mature populations co-transferred in bulk bone marrow after BCG administration, is capable of conferring a systemic anti-tumor effect.

A recent study observed recruitment of pro-tumorigenic immunosuppressive CCL3^hi^ PDL1^hi^ neutrophils by IL-8 secreted by bladder cancer cells (62), consistent with another recent report of pro-angiogenic tumor supporting neutrophils (63). We asked if BCG administration altered this pro-tumor neutrophil phenotype and function. Consistent with BCG-induced changes in neutrophil phenotypes, bladder administration of BCG shifted the density of bone marrow progenitor cells from the monocyte lineage to the granulocyte lineage (Figure 3C). This observation is consistent with data from intradermal BCG vaccination, which induces increased granulopoiesis and functional reprogramming of neutrophils with increased antimicrobial capacity (64). To understand the contribution of neutrophils to the tumor control conferred by BCG reprogrammed HSCs, we generated chimeric animals from either PBS or Bladder BCG donor mice, challenged with subcutaneous MB49, and depleted neutrophils with anti-Ly6G depleting antibody (Figure S4D). Similar to previously published results(62), tumors were significantly smaller when neutrophils were depleted in mice that received control bone marrow, consistent with a pro-tumorigenic role for neutrophils. However, the enhanced tumor control conferred by BCG bone marrow was lost when neutrophils were depleted, indicating that BCG converts neutrophils away from a pro-tumor phenotype. Seminal studies by Lloyd Old and colleagues identified TNFɑ as the principal serum factor induced by BCG that confers resistance to tumor challenge (65). To investigate the role of TNFɑ in conferring HSPC transplantable tumor control, we treated chimeric animals with a TNFɑ with blocking antibody and observed that a loss of the tumor control conferred by BCG-experienced HSPC (Figure S4E). These results indicate that transplantation of BCG-experienced HSPC contributes to improved control of tumor growth through mechanisms involving both altered neutrophil phenotypes and the production of TNF by HSPC-derived immune cells.

### BCG-reprogrammed hematopoietic stem and progenitor cells confer enhanced tumor infiltration to mature innate immune cells

To characterize the effect of BCG-induced innate immune memory in the bladder tumor microenvironment (TME), we generated groups of congenically-marked mixed bone marrow chimeras according to the schematic shown in Figure 5, wherein bone marrow from CD45.2^+/+^ donor mice treated with either bladder or intravenous BCG was mixed 1:1 with naive CD45.1^+/-^CD45.2^+/-^ bone marrow. The 1:1 mixed donor bone marrow in all combinations was used to reconstitute naive irradiated CD45.1^+/+^ recipient mice (Figure 5A).

**Figure 5:**
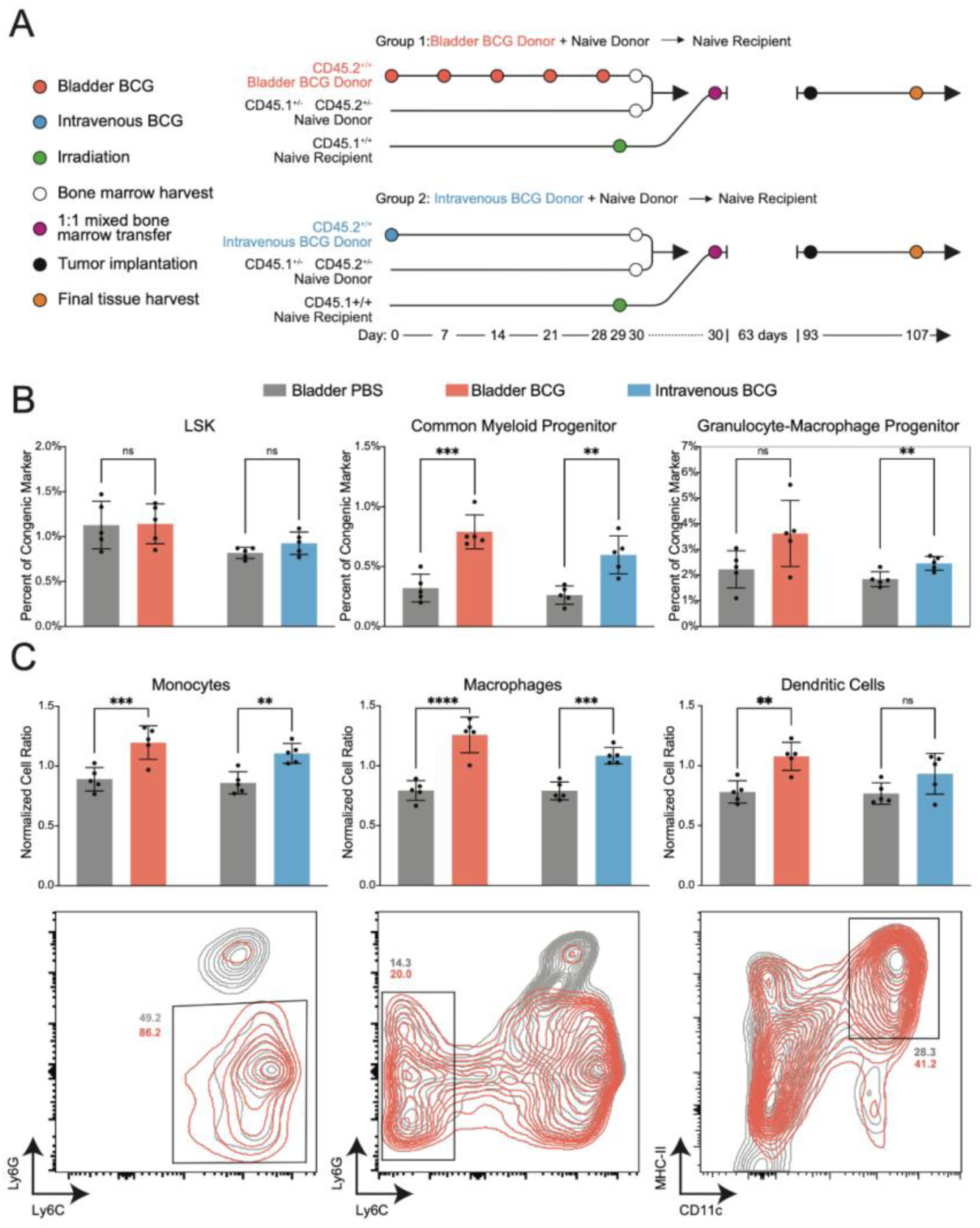
BCG-experienced bone marrow confers enhanced tumor infiltration to mature innate immune cell progeny. A) Experimental schematic. Mixed bone marrow chimeras were generated by transferring a 1:1 mix of bone marrow from CD45.2^+/+^ donors treated with 5 weekly doses of bladder BCG and CD45.1^+/-^ CD45.2^+/-^ naive donors (Group 1), or CD45.2^+/+^ donors treated with single-dose intravenous BCG and CD45.1^+/-^ CD45.2^+/-^ naive donors (Group 2), into naive irradiated CD45.1^+/+^ recipient mice. After immune reconstitution from donor bone marrow, recipient mice were challenged with MB49 bladder tumors. Tumors, spleens, and bone marrow were harvested 2 weeks post-implantation and analyzed by flow cytometry. B) Quantification of bone marrow HSPC populations from bladder or intravenous BCG-experienced origin *versus* naive origin in Group 1 and Group 2, respectively. C) Quantification of tumor-infiltrating myeloid cell populations from bladder or intravenous BCG-experienced origin *versus* naive origin in Group 1 and Group 2, respectively (top). Values represent fold change of cell frequency in the tumor compared to cell frequency in the spleen within each congenically-marked cell type. Representative flow plots are shown (bottom). Percentages shown represent the frequency of the parent gate within each congenic marker. Monocytes (left) were gated by CD45 congenic marker, CD11b+, F4/80-, Ly6G-, and Ly6C+. Macrophages (middle) were gated by CD45 congenic marker, CD11b+, F4/80+, Ly6G-, and Ly6C-. Dendritic cells were gated by CD45 congenic marker, F4/80-, CD11c+, and MHC-II+. P values were derived by Student’s t-test. P > 0.05 = ns, P ≤ 0.05 = *, P ≤ 0.01 = **, P ≤ 0.001 = ***, P ≤ 0.0001 = ****.

After engraftment, we confirmed greater than 95% immune reconstitution from mixed donor bone marrow in all mice by flow cytometry, ensuring that subsequent hematopoiesis reflected the engrafted donor HSPCs. Analysis of the bone marrow revealed similar proportions of LSKs between the two donors in all three groups (Figure 5B). However, we observed preferential myeloid progenitor expansion originating from bladder and intravenous BCG-experienced bone marrow origin compared with cells of the naive donor origin, consistent with our prior findings of BCG-induced myelopoiesis (Figure 5B, S5A). Specifically, common myeloid progenitor (CMP), common monocyte progenitor (cMoP), granulocyte-monocyte progenitor (GMP), and neutrophil progenitor (NP) originating from BCG-experienced progenitors were all more abundant, with a similar magnitude of enrichment between the two routes of BCG administration (Figure 5B and Figure S5).

These chimeric animals were then challenged with bladder tumors to determine competitive cell-intrinsic phenotypes within the TME. Because we observed myeloid skewing in the BCG-experienced HSPC (Figure 5B, S5A) we analyzed the bladder infiltration of immune cells by normalizing to the spleen to control for bone marrow output and estimate cell intrinsic differences in tumor migration and proliferation. Tumors implanted in these chimeric mice revealed that mature tumor-infiltrating myeloid cells originating from BCG-reprogrammed HSPCs were preferentially enriched compared to tumor-infiltrating cells of naive HSPC origin. Specifically, we found an increased relative abundance of monocytes, macrophages, and DCs from both the bladder and intravenous BCG-experienced donor groups (Figure 5C). In contrast, analysis of the bulk T cell populations revealed no significant differences in abundance of T cell subsets of naive or BCG-experienced origins in either group (Figure S5B-D), indicating that the effects of BCG on HSPCs are not conveyed to T cell progeny in a manner that augments their anti-tumor activity in a direct and cell-intrinsic manner. Together, these data indicate that reprogramming of HSPCs by BCG confers an augmented cell-intrinsic capacity for tumor infiltration to progeny myeloid cells.

### HSPC-reprogramming by bladder BCG augments T cell priming and synergizes with immune checkpoint inhibitor therapy to enhance tumor clearance

Our results establish that administration of BCG into the bladder has a systemic effect, and cells derived from BCG-experienced HSPCs, particularly dendritic cells and macrophages, have enhanced antigen presentation pathways and preferentially migrate to the tumor microenvironment. Although we did not detect an effect of BCG-reprogrammed HSPCs on bulk tumor T cell numbers (Figure S5), we tested whether tumor specific T cell responses were enhanced by BCG-reprogrammed HSPCs. Bone marrow chimeric mice reconstituted with LSKs from bladder PBS-, bladder BCG-, and intravenous BCG-experienced donors were implanted with MB49 tumors expressing both MHC Class I (MHC-I) and Class II (MHC-II) epitopes of the model neoantigen ovalbumin (OVA), as depicted in Figure 6A. Seven days after tumor implantation, mice received congenically-marked OT-I (CD8) and OT-II (CD4) transgenic T cells specific to the MHC-I and MHC-II epitopes of OVA, respectively. Transferring naive transgenic T cells allowed us to ensure that any changes in T cell phenotype would arise *via* the observed epigenetic and transcriptional enhancements to the myeloid compartment. Five days after the T cell transfer, bladder tumors were harvested for analysis by flow cytometry (Figure 6A). We observed an increased infiltration of OVA-specific CD8 T cells, but not OVA-specific CD4 T cells, in both bladder and intravenous BCG- experienced bone marrow chimeras *versus* the control group (Figure 6B).

**Figure 6:**
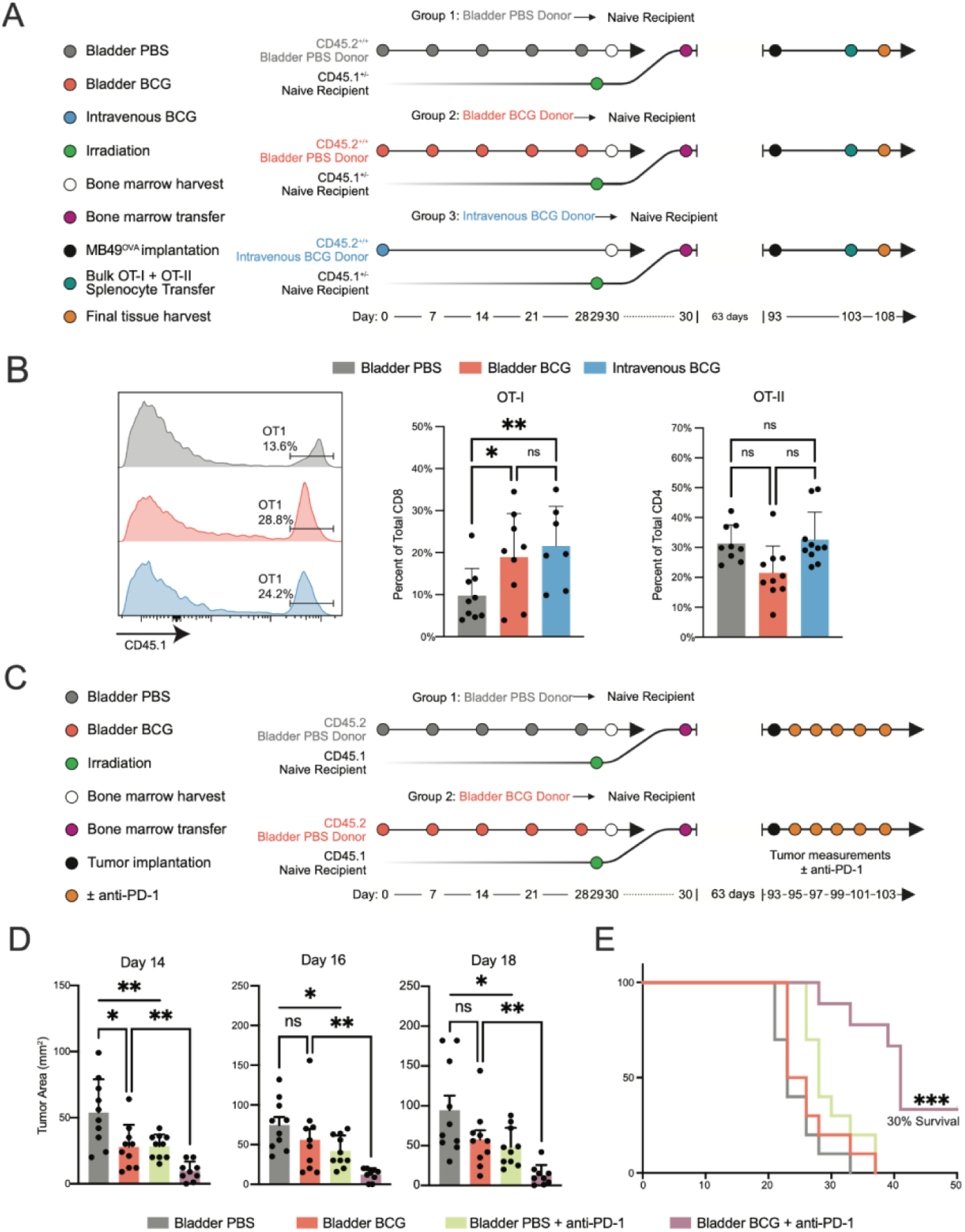
BCG-induced HSPC-reprogramming enhances the tumor-specific T cell response and synergizes with checkpoint blockade. A) Experimental Schematic. Bone marrow chimeras were generated by transferring bulk bone marrow from CD45.2^+/+^ donor mice treated with 5 weekly doses of bladder BCG, 5 weekly doses of bladder PBS, or single-dose intravenous BCG, into naive irradiated CD45.1^+/-^ recipient mice. After 9 weeks to allow for full reconstitution of the immune system from donor bone marrow, chimeric mice were challenged with bladder MB49^OVA^ tumors, and CD45.1^+/+^ OT-I and OT-II T cells 10 days later. Bladder tumors were harvested 5 days after T cell transfer to assess tumor-specific T cell frequency. B) Representative histograms depicting the frequency of OT-I T cells in bladder tumors among total CD8+ cells in bladder PBS-, bladder BCG-, and intravenous BCG- experienced bone marrow recipients are shown at left. Quantification of OT-I and OT-II T cell frequency for all groups is shown at right. C) Experimental Schematic. Bone marrow chimeras were generated by transferring bulk bone marrow from CD45.2^+/+^ donor mice treated with 5 weekly doses of bladder PBS or bladder BCG into naive irradiated CD45.1^+/+^ recipient mice. After reconstitution, chimeric mice were challenged with either subcutaneous or bladder MB49 tumors to assess tumor growth and survival, respectively. Mice in both groups were administered 5 doses of anti-PD-1 or PBS every 2 days following tumor implantation to assess the synergistic effects of trained immunity with checkpoint blockade. D) Subcutaneous MB49 tumor growth comparisons for Days 14, 16, and 18 from the experiment described in C. E) Survival following MB49 bladder tumor challenge from the experiment described in C. P values for bar graphs were derived by Student’s t-test. P values for survival curves were derived by log-rank test. P > 0.05 = ns, P ≤ 0.05 = *, P ≤ 0.01 = **, P ≤ 0.001 = ***, P ≤ 0.0001 = ****.

Despite the enhanced tumor control and anti-tumor immune responses conferred by BCG-exposed HSPCs, MB49 tumors always progressed, suggesting that HSPCs alone are insufficient to induce a complete and protective anti-tumor immune response. We hypothesized that BCG-induced innate immune memory may synergize with T cell-targeting immune checkpoint inhibitor therapy. To test this hypothesis, we challenged bone marrow chimeric mice reconstituted with BCG-experienced or control HSPCs with subcutaneous MB49 tumors and treated them with anti-PD-1 (Figure 6C). As previously shown, mice that were reconstituted with BCG-experienced bone marrow demonstrated enhanced control of tumors over the PBS control group (Figure 6D). PBS control bone marrow chimeras treated with anti-PD-1 demonstrated a similar level of tumor control to the BCG-experienced bone marrow chimeras (Figure 6D). However, the combination of BCG-experienced HSPCs and anti-PD-1 exhibited significant reductions in tumor volume and approximately 30% survival, demonstrating a strong synergistic effect of BCG-induced innate immune memory and checkpoint blockade (Figure 6D-E, Figure S5D).

## Discussion

The development of BCG as the first cancer immunotherapy emerged from studies of the effect of microbes and microbial products on tumor growth (66–68). Despite its well-established clinical efficacy, the detailed mechanisms by which BCG controls tumors have remained elusive. Recent evidence in mouse models and human patients indicated that BCG-mediated tumor rejection is dependent on tumor specific T cells (21–23). The accepted model of anti-tumor effects of BCG, and the rationale for its administration in the bladder, was that it acts as a local immunotherapy at the site of administration to improve anti-tumor T cell priming. Upstream immunologic events stimulated by BCG that enable this immunity were unknown.

Although the fundamental principle of vaccination is the induction of antigen-specific T and B cell immunity, it has become clear that BCG vaccination elicits innate immune responses that can cross-protect against antigenically unrelated pathogens, including viruses (69–74). A prominent, recently appreciated embodiment of this phenomenon, termed innate immune memory or trained immunity, is the reprogramming of hematopoietic progenitors in the bone marrow and subsequent skewing of the abundance and function of myeloid cell progeny (27–29). It has been unknown if this progenitor encoded or “central” innate immune memory contributes to the anti-tumor effects of BCG and if administration of BCG directly into the bladder could induce central innate immune memory.

Our data revises the understanding of BCG’s anti-tumor effects, including in the context of its longstanding clinical use as the first microbial immunotherapy, demonstrating potent systemic rather than only local effects. Bladder administration modifies the epigenetic and transcriptional landscape of bone marrow hematopoietic stem cells. BCG administered to the bladder mucosal epithelium in mice traffics to the bone marrow, a finding we directly demonstrate by cultivating viable BCG from the bone marrow. These data indicate that the hematopoietic-reprogramming previously implicated in the heterologous protection from infection conferred by early life intradermal BCG vaccination in humans (26,33,71) and by intravenous administration in mice (27, 39, 75) is a shared and intrinsic part of BCG-mediated immunotherapy of cancer. Furthermore, BCG-reprogrammed HSPCs were sufficient to confer anti-tumor immunity to recipient mice, indicating a durable and persistent cell-intrinsic memory in progenitor cells that is conveyed through differentiation to mature myeloid cells. Our single-cell ATAC and RNA sequencing data from mice and humans, coupled with functional characterization of mixed bone marrow chimeras, indicate that a broad enhancement of myeloid function contributes to the antitumor effects of innate immune memory. These include a functional dependence on neutrophils and TNF, as well as enhanced infiltration of tumors with macrophages and DCs that encode enhanced antigen presentation pathways. These HSPC programs include epigenetic and transcriptional changes that are conveyed through differentiation to mature myeloid cells. Importantly, we observed increased abundance of DC progeny from BCG- experienced HSPCs in tumors, along with enhanced numbers of tumor-specific T cells, suggesting that our previous observations of augmented tumor infiltrating T-cells are driven by BCG-induced innate immune activation (21). Of substantial clinical significance, the myeloid cell progeny of BCG-experienced HSPCs strongly amplify the response to PD-1 blockade, thereby directly coupling innate immune memory to the anti-tumor T cell response.

Our findings have implications for the use of BCG in bladder cancer but also as a broad immunotherapy against other cancers. BCG remains the standard of care for non-muscle-invasive bladder cancers, but a substantial minority of treated patients will experience tumor recurrence and there are no reliable pre-treatment predictors of response (76). Our data suggest that inter-individual differences in either pre-treatment or BCG-induced innate immune memory could predict the response to BCG, a hypothesis that can be tested using the PBMC-PIE approach (43). More broadly, our data suggest that augmenting the abundance and function of the myeloid compartment through HSPC-reprogramming is an effective method to improve anti-tumor immunity, including and especially in combination with immune checkpoint blockade. These concepts are supported by a recent study that revealed a pro-tumor hematopoietic circuit involving IL-4 that could be therapeutically targeted to improve anti-tumor responses (77), and another that described a bone marrow targeting peptidoglycan that induces HSPC reprogramming, increased myelopoiesis, and improved tumor control (36). There is substantial evidence that dendritic cell abundance and function are important determinants of checkpoint blockade activity (78–82), but harnessing this knowledge for DC-derived therapies is limited by the short lifespan of these cells. Our data suggests that HSPC-reprogramming, including by BCG administered to the bladder, may provide a strategy for more durable alterations in myeloid function to enable successful anti-tumor immune responses systemically and across a range of anatomical locations and tumor types.

## Acknowledgments

MSKCC FCCF Core for assistance with panel generation and troubleshooting. Rachel E. Niec (MSKCC) for extensive discussions and helpful suggestions on the science and manuscript.

We thank the Josefowicz and Glickman Labs for discussions and paper review Immunology and Microbial Pathogenesis Programs for feedback and support

## Funding

Department of Defense Horizon Award CA181350 (ACA)

National Institutes of Health grant 5F31HL152706 (AWD)

National Institutes of Health grant 5T32AI134632 (ACA, AWD)

Ludwig Center at Memorial Sloan Kettering Cancer Center (MG, GRS)

National Institutes of Health grant P50CA221745 (MG, GRS)

National Institutes of Health support grant P30 CA008748 (MG, GRS)

National Institutes of Health grant R01AI148416 (SZJ)

National Institutes of Health grant R01AI148416-S2 (SZJ)

Burroughs Wellcome Fund Pathology Award (SZJ)

Hirschl Weill-Caulier Award (SZJ)

Bochner-Fleisher Research Grant (GRS)

Canadian Institute of Health Research Project Grant MM1-174910 (MD)

Fonds de Recherche du Québec-Santé Award (MD)

Strauss Chair in Respiratory Diseases (MD)

Fellow member of the Royal Society of Canada (MD)

Fonds de Recherche du Québec—Santé studentship (LFJ)

## Author contributions

Conceptualization: ACA, AWD, MSG, SZJ Data curation: LP

Formal Analysis: ACA, AWD, GRS, JGC, MJB, LP

Funding acquisition: MSG, SZJ, ACA, AWD, MD

Investigation: ACA, AWD, GRS, LFJ, DJA, SK, MD, OL, VAM, AB, SJ, EP

Methodology: ACA, AWD, MSG, SZJ

Supervision: MSG, SZJ

Validation: ACA, AWD, GRS

Visualization: ACA, AWD, JGC, MJB, LP

Writing – original draft: ACA, AWD, MSG, SZJ

Writing – review & editing: ACA, AWD, MSG, SZJ, LFJ, DP, MD

## Data and materials availability

Sequencing data and analysis code will be publicly available in Zenodo repository: https://doi.org/10.5281/zenodo.10695064. All other data are available in the main text or the supplementary materials.

## Supplementary Information

Figures S1-S5

Tables S1-S2

## Materials and Methods

### Cell lines

The mouse bladder cancer cell line MB49, expressing luciferase under G418 selection, was a gift from Yi Luo, University of Iowa, Iowa City, IA. The mouse melanoma cell line B16 was obtained from Taha Merghoub, Memorial Sloan Kettering Cancer Center, New York, NY. MB49 and B16 were grown in RPMI supplemented with 10% FBS, and 2 mM L-glutamine. Cells were cultured at 37 °C in a humidified atmosphere of 5% CO2. All cell lines used were confirmed to be negative for mycoplasma by annual testing using MycoAlert Plus (Lonza). Last testing was performed on 14 August 2019.

### BCG

The Pasteur strain of BCG was grown at 37℃ in Middlebrook 7H9 supplemented with 10% albumin/dextrose/saline, 0.5% glycerol, and 0.05% Tween 80. To create titered stocks for infection, BCG was grown to mid-log phase (OD600 0.4 to 0.6), washed twice in PBS with 0.05 Tween 80, resuspended in PBS with 25% glycerol, aliquoted, and stored at -80℃. To measure the final bacterial titer, an aliquot was thawed and serial dilutions were cultured on 7H10 agar. The bacterial titer was determined by counting colonies after 3 weeks of incubation.

### Mouse strains

Wild-Type C57BL/6 (Strain #: 000664), CD45.1 (Strain #: 002014), OT-I (Strain #: 003831), and OT-II (Strain #: 004194) mice were purchased from The Jackson Laboratory. All mouse strains were bred and housed in Memorial Sloan Kettering Cancer Centers (MSKCC) Research Animal Resource Center under specific pathogen-free conditions. All animal studies were performed with approval from the MSKCC Institutional Animal Care and Use Committee Under Protocol 01-11-030 and were compliant with all applicable provisions established by the Animal Welfare Act and the Public Health Services Policy on the Human Care and Use of Laboratory Animals.

### MB49 orthotopic implantation

Seven- to eight-week-old female mice (The Jackson Laboratory) were placed under anesthesia in an isoflurane chamber. Mice were transferred from the chamber to a nose cone for the procedure and returned to the chamber for incubation steps. For each mouse, a 24-gauge catheter (Terumo) was inserted into the bladder through the urethra. Next, 100 μL of poly-L-lysine (Sigma) was injected through the catheter, the catheter was capped using an injection plug (Terumo), and the mice were kept under anesthesia for 30 minutes. After 30 minutes, catheters were removed from one mouse at a time in the same order as they were implanted. The catheter was then flushed with a solution containing 500,000 MB49 cells/mL in RPMI. Each mouse was then removed from the isoflurane chamber in turn, the bladder was manually emptied, and the catheter was re-inserted. 100 μL of the MB49 solution (50,000 cells/mouse, unless otherwise noted) was injected into the bladder and the catheter was re-capped. The mice were kept under anesthesia for 1 additional hour. At the end of the hour, catheters were removed and the mice were allowed to recover from anesthesia. Mice were observed daily and were euthanized if they displayed signs of distress, such as dull fur, apathy, or visible signs of growing tumor.

### BCG administration

Frozen titered stocks of BCG (prepared as detailed above) were thawed and resuspended in PBS for a final concentration of 3x10^7^ colony forming units/mL (CFU/mL). PBS alone was used as a control. Mice were placed under anesthesia in an isoflurane chamber, transferred from the chamber to a nose cone for the procedure, and returned to the chamber for incubation steps. For intravesical administration, a 24-gauge catheter (Terumo) was inserted into the bladder through the urethra, 100 μL of BCG (3x10^6^ CFU/mouse) was injected into the bladder, and the catheter was capped using an injection plug (Terumo). The mice were kept under anesthesia for 2 hours, after which catheters were removed and mice were allowed to recover from anesthesia. For intravenous BCG (administered via retro-orbital injection), mice were placed in a left lateral position, and gentle pressure was applied above and below the eye to protrude the ocular globe. A 30-gauge needle was carefully inserted approximately 2 mm into the posterior eye socket, and 100 μL of BCG (3x10^6^ CFU/mouse) was injected into the retro-orbital sinus. The needle was removed and the mice were allowed to recover from anesthesia.

### Measurement of subcutaneous tumors

Subcutaneous tumor measurements were obtained using a caliper by measuring the longest axis of the tumor first, followed by the perpendicular axis.

### Flow cytometry

Cell suspensions were analyzed on a LSR Fortessa (BD Biosciences) or Aurora (Cytek), using FACS DiVa software (BD Biosciences) or SpectroFlo (Cytek), respectively. Data analysis was performed using the FlowJo software package (Tree Star). For determination of cytokine production by T cells, single cell suspensions were restimulated with 1X Cell Stimulation Cocktail (plus protein transport inhibitors) (eBioscience) for 6 h at 37℃. Cells were first stained with a fixable viability dye, followed by surface markers, then fixed and permeabilized using Foxp3 Fixation/Permeabilization Buffer (eBioscience) according to the manufacturer’s instructions, and finally stained for intracellular antigens. Antibodies used for this study are detailed in Table 2.

### Bone marrow chimeras

Whole body irradiation using a cesium source was used to induce hematopoietic ablation of live recipient mice. Mice were placed in a rotating pie-shaped holder then placed in the machine where they received a 9 Gy dose of radiation. Each irradiated mouse received 10^6^ to 10^7^ donor cells (whole bone marrow or sorted LSK cells, as appropriate) in 0.1 mL sterile PBS via retro-orbital injection within 24 hours post-irradiation. Mice were monitored 3 to 4 times per week for 2 weeks post-irradiation to ensure no acute illness occurred. All recipient mice were rested for a minimum of 8 weeks to allow full immune reconstitution before further experimentation as described throughout this manuscript. After 8 weeks, a small amount of blood was drawn to confirm full reconstitution of the immune system from donor cells by flow cytometric analysis of immune cell lineages.

### T Cell Isolation and Adoptive Transfer

Donor spleens were harvested and single-cell suspensions were made. CD4 and CD8 T cells were isolated using mouse T Cell Isolation Kits (Miltenyi). After cells were counted, recipient mice were placed under anesthesia in an isoflurane chamber and 3 to 5 million cells per mouse were transferred via retro-orbital injection.

### PBMC-PIE

For each human sample, we prepared two conical tubes with RPMI (labeled as tube1 and tube2). We thawed frozen PBMCs in a 37°C water bath and transferred them into tube1. Subsequently, 10% of this suspension was transferred to tube2 for genotyping and sorting viable PBMCs into enriched CD34+ HSPC. Both tubes were centrifuged at 300g for 10 minutes. The resulting pellets were resuspended in MACS buffer. We combined the pellets from 6-8 samples into one tube and proceeded with CD34+ cell enrichment using CD34 microbeads (Miltenyi #: 130-046-702). To optimize the yield of HSPC, the MACS column was not washed, the CD34- fraction (flowthrough) was re-added to the column and then cells were removed from the magnet and eluted. The enriched cells collected in the conical tube were then centrifuged and resuspended in FACS staining buffer containing the following antibodies: FITC anti-CD34 (Miltenyi #: 130-113-178, 1:100), Pacific Blue anti-CD49f (Biolegend #: 313620, 1:200), PE anti-CD90 (Biolegend #: 328110, 1:100), PE-Cy7 anti-CD38 (Biolegend #: 303516, 1:100), APC-Cy7 anti-CD45RA (Biolegend #: 304128, 1:400), and anti-lineage (cat number). Staining was performed for 30 minutes in the dark. Post-staining, cells in both tube1 (CD34+ enriched cells) and tube2 were resuspended in 7-AAD-containing MACS buffer for sorting. Initially, viable lineage-negative cells were sorted into a PCR tube, followed by sorting PBMCs from tube2 into the same PCR tube. The number of PBMCs sorted was determined based on the desired ratio of PBMC and CD34+ cells in the data.

### Mouse progenitor enrichment

For bone marrow isolations, we harvested bone marrow from tibia and femurs, after RBC lysis, cells were stained for lineage markers (CD3, NK1.1, Gr-1, B220, Ter119), c-Kit and Sca-1. 90k viable, lineage-negative cells were sorted into a PCR tube, and then 10k of lineage positive cells were sorted into the same PCR tube, allowing for a small representation of lineage positive cells in our dataset.

### 10X Multiome

Following sorting, we immediately proceeded with the 10X Multiome protocol. Nuclei isolation was conducted following the low-yield nuclei procedures in the appendix of the nuclei isolation protocol provided by 10X Genomics. The rest of the steps were performed as per the manufacturer’s manual. Sequencing libraries for ATAC-seq and RNA-seq were generated and sequenced using Novaseq6000.

### Preprocessing of single-cell multiome sequenced data

The Cell Ranger ARC 2.0.2 pipeline was used for initial processing (sample demultiplexing, barcode processing, alignment of reads, counting of transcripts, cell filtering) of all human and mouse single-cell multiome data with the hg38 and mm10 reference genome.

### Mouse Antibody Administration

For depleting or blocking experiments (TNF(BE0058) or Ly6G(BE00775-1) mice were treated 2 days before tumor cell challenge, and then every 2-3 days subsequently (Days: -2, 0, 5, 7,9) with 200ug antibody per mouse administered IP.

### Human single-cell multiome data processing and basic analysis

Starting from initial Cell Ranger filtered cells, we performed additional manual filtering per sample to ensure only high-quality cells remained in our data: iteratively embedding and clustering the data, removing clusters with poor quality-control (QC) metrics, and then re-embedding and clustering. RNA data was processed using Scanpy 1.9.3 (83) (median and log normalization of counts, PCA, and UMAP), and ATAC data was processed using ArchR 1.0.1 (84) (iterative LSI, UMAP). Clustering was run on the respective PCA and LSI matrices using PhenoGraph (85). Cluster QC metrics evaluated included standard Scanpy and ArchR-calculated metrics, DoubletDetection score, and mitochondrial and ribosomal fraction.

Data from separate samples was integrated without additional data harmonization. For the RNA modality, count matrices from each sample were concatenated into a full data matrix, which was then median and log-normalized. Ribosomal and mitochondrial genes were removed. The top 45 PCs were calculated using 1500 highly variable genes (HVG). We ran PhenoGraph clustering and UMAP using 30 nearest neighbors. For the ATAC modality, we applied ArchR’s implementation of iterative LSI to the tile matrix using 100,000 variable features, and ran PhenoGraph on the 30 nearest neighbors in LSI embedding coordinates. Cells were annotated based on manual evaluation of PBMC marker gene expression in RNA clusters. These steps were performed first on cohort 1 and cohort 2 samples independently, and then on all samples combined.

### Entropy analysis for the evaluation of single-cell multiome batch effect

We calculated entropy as described by (86). We first constructed a k-NN(k=30) graph in RNA PCA space using Euclidean distance. Given a cell *i* and its 30 nearest neighbors, we computed the fraction of cells from each sample s = 1,…,10 as *p_i_^s^*. We find the Shannon entropy per cell as:

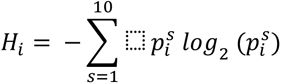

High entropy indicates that a cell’s neighborhood in RNA space is made up of a well-mixed set of samples, whereas low entropy indicates that nearby cells mostly come from the same sample.

### Human single-cell multiome differential gene expression

To determine differentially expressed genes post-BCG treatment, we employed MAST, a hurdle model that accounts for the many zero-counts in scRNA-seq data (87). For each cell type, we fit the MAST model to log-normalized RNA counts of post vs. pretreatment cells, returning a false discovery rate (fdr) and MAST coefficient (Coef.), per gene per cell type. MAST Coef. can be interpreted similarly to log fold change, it indicates the direction and magnitude of change in gene expression between groups.

Due to an overrepresentation of female cells in our HSPC cluster, we found significant genes in MAST results for that cluster to be dominated by sex-linked genes. For HSPC, we ran MAST a second time excluding the genes XIST, TSIX, and all genes on the Y-chromosome. MAST results including all genes for HSPC can be found in supplemental table 1.

Gene set enrichment analysis was performed on notable gene sets throughout this project using EnrichR (88), comparing a given set of genes to the GO Biological Process 2021 reference (89,90).

### Chromvar motif accessibility

We constructed a reproducible peak set for each iteration of our human multiome dataset (cohort 1, cohort 2, and combined) using ArchR, grouping cells by ATAC PhenoGraph clusters before calling and merging peaks. We annotated motifs within peaks using the CISBP motif database, and determined chromVAR score per cell using ArchR’s addBgdPeaks() and addDeviationsMatrix() functions.

To evaluate changes in motif accessibility post-treatment, we compared chromVAR scores in all post-treatment vs. pre-treatment cells for each motif and cell type using a Wilcoxon rank-sum test. Statistical significance of motifs was decided based on an p- value cutoff of 0.05. We also calculated the mean difference (MeanDiff) in chromVAR score post v. pre-treatment as the mean of cell scores for a given cell type and motif pre-treatment subtracted from the mean of scores post-treatment.

### Mouse Single-Cell Multiome Data Processing and Basic Analysis

For the analysis of mouse single-nuclei multiome datasets, we employed R packages Seurat and Signac (91,92). We initiated the process by utilizing Cell Ranger outputs for filtered cells to create Seurat objects for each sample. Subsequently, we conducted manual cell filtering for each sample, eliminating cells with either excessively high or low fragment numbers or RNA counts per cell, along with cells exhibiting low TSS enrichment scores. We filtered out cells with RNA count less than 100, ATAC fragment count less than 1000, or cells with unusually high counts.

Next, we utilized the ATAC-seq assay within the Seurat object to call peaks for each sample using MACS2. These peaks were then combined using the ’reduce’ function of GenomicRanges. Following this step, we generated peak count matrices once again for each sample and created a merged Seurat object. We then applied the standard Signac workflow, including TF-IDF normalization and SVD with default parameters. UMAP embeddings were generated from the first 50 dimensions obtained through the LSI reduction method. Finally, we computed nearest neighbors using the default settings of the Signac package.

Utilizing the RNA-seq assay of Seurat objects for each sample, we created another merged Seurat object, which was then divided into layers by sample using the ’split’ function. Standard Seurat preprocessing workflow steps such as normalization, scaling, and PCA were carried out. The split layers were integrated using ’integrateLayers,’ resulting in a new dimensional reduction labeled ’integrated.cca.’ The layers were subsequently rejoined using the ’JoinLayers’ function within the Seurat package. We did not further process the RNA-seq dataset for UMAP embedding and clustering analysis.

For motif analysis, we incorporated motif information into the merged object using the ’AddMotifs’ function in Signac. Motif information for mm10 was obtained from the JASPAR2020 database. Additionally, we added per-cell TF motif activity scores using chromVAR with the ’RunChromVAR’ function in Signac as a separate assay to the object.

Due to the limited depth of the RNA-seq dataset, we were unable to derive meaningful clusters based on transcriptome data. Consequently, we relied on the cluster information derived from the ATAC-seq assay to identify cell types. When annotating each cluster, we referenced the cell type calling results from SingleR package (93) , and with the expression of cluster marker genes and major cell type-specific chromVAR TF activity.

To analyze differential TF activity (chromVAR scores) across groups, we utilized the ’FindMarkers’ function in Seurat, applying the Wilcoxon test. For differential gene expression analysis, the Wilcoxon test was also employed, enabling us to generate volcano plots that highlight significantly differentially expressed genes with adjusted p- values less than 0.05. To correlate differential gene expression with differential chromVAR TF activity, we employed the MAST (92) test to identify genes that were significantly differentially expressed.

### Human-mouse differential gene expression comparison

Human-mouse orthologous genes were matched based on the HGNC Comparison of Orthology Predictions (HCOP) search tool. Not all matches were one-to-one; a number of mouse genes in our data were mapped to multiple human orthologs. For each gene-gene pair, we compared differential expression results from mouse and human single-cell multiome datasets: specifically, comparing MAST Coef. in humans to Log2FC in mouse.

### Bulk ATAC sequencing of mouse LSKs

For ATAC-seq, we followed the Omni- ATAC-seq protocol, working with 50,000 LSK cells sorted directly into PCR tubes.

### Bulk ATAC-seq Data Processing

ATAC-seq fastq files were processed using an in-house pipeline implemented in nextflow () at https://github.com/michaelbale/jlabflow. Briefly, paired-end reads were trimmed for low-quality base-calls and adapter contamination using the Cutadapt () wrapper Trim Galore (). Remaining reads were then mapped to mm10 using Bowtie2 () with parameters “`--no-mixed --no-unal --no-discordant –-local -–very- sensitive-local -X 1000 -k 4 --mm”` retaining only properly mapped fragments with a MAPQ score of at least 30. Mitochondrial reads and improperly paired reads or secondary alignments were removed with Samtools () and Picard () was used to remove duplicate fragments. Finally, we removed all mapped fragments that were associated with the ENCODE Forbidden list ().

### Genome Browsing Tracks

Genome browsing tracks were generated as bigwig files using Deeptools bamCoverage () with reads per genomic content normalization using an effective genome size of 2648000000. For figures XXX, bigwigs from individual replicates were averaged together using Deeptools bigwigAverage.

### Analysis of ATAC-Seq Data

Peak calls for individual samples were made using Genrich () in ATACseq mode (“-j”). Reproducible peaks within each treatment condition were determined using ChIP-r () and optimal peak calls between conditions were merged to form an atlas of 25,245 total peaks. Reads in peaks were generated by Deeptools multiBamSummary and read in to R v4.3.0 for differential analysis using DESeq2 v1.40.2 (). Finally, motif bias analysis was performed using HOMER2 findMotifsGenome.pl () with input peaks as peaks that were differentially accessible in BCG-treated LSK over PBS-treated (as defined by DESeq2 analysis) using differentially accessible peaks in PBS-treated LSK over BCG-treated as the custom background set (-bg).

---

**Figure S1:**
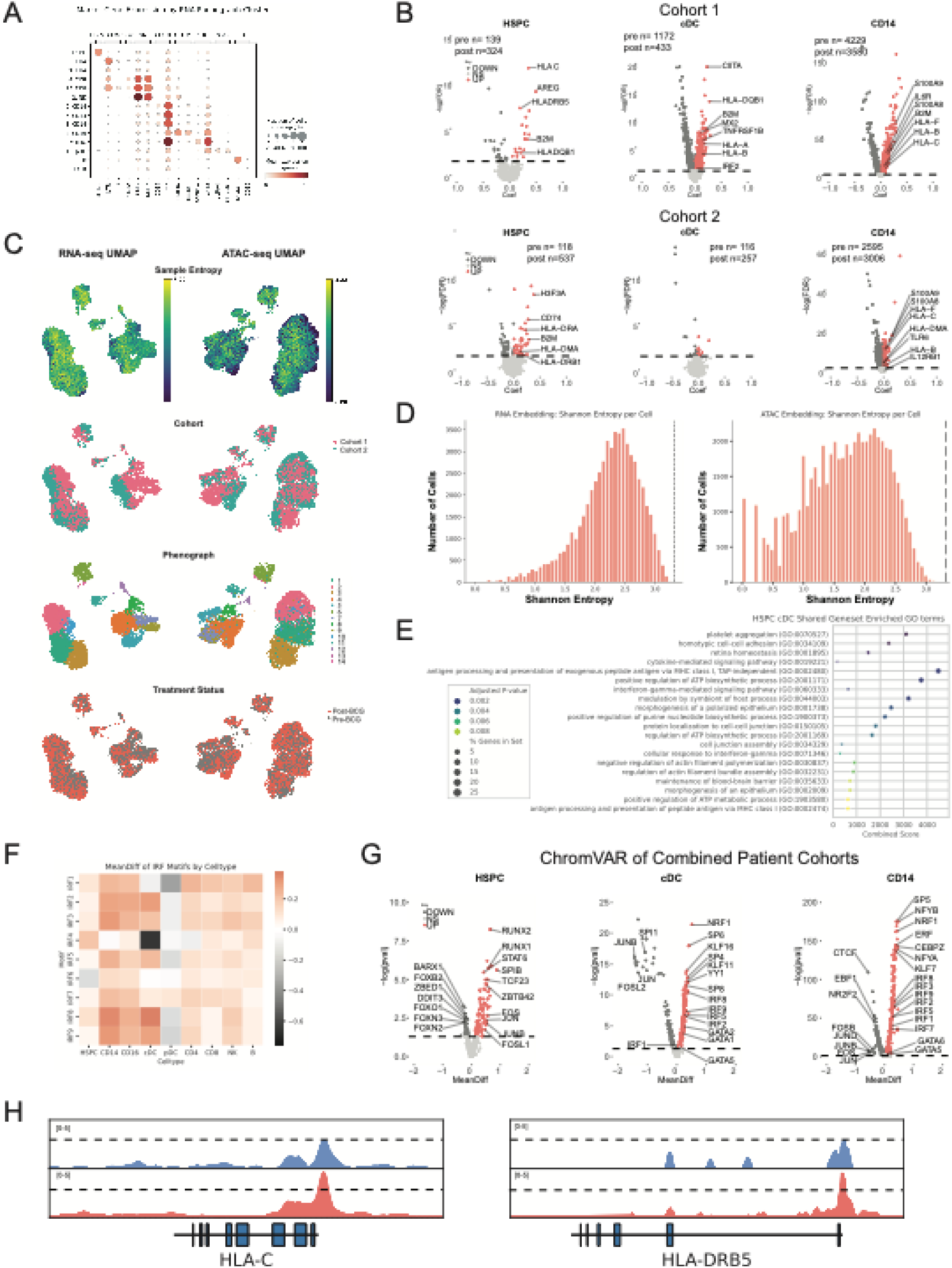
Multi-omic analysis reveals trained HSPC phenotypes in BCG-treated bladder cancer patients at two separate institutions. **A**) Dot plot of marker gene expression (columns) in each RNA PhenoGraph cluster (rows). Clusters are labeled by cluster number and the corresponding annotated cell type association. Dots are colored by mean expression per cluster, and dot size corresponds to the fraction of cells in the cluster with non-zero expression of that gene. **B)** Volcano plots of differentially expressed genes post- *versus* pre-BCG in HSPCs, cDCs, and CD14+ monocytes in Cohort 1(top, MSKCC) or Cohort 2 (bottom, McGill). Significant differentially expressed genes (fdr < 0.05) are colored according to their relative enrichment up or down, and the number of cells analyzed per group is given above each plot. **C)** UMAP visualizations of RNA and ATAC data modalities from all collected samples, colored by (top to bottom) sample entropy score, patient cohort, RNA PhenoGraph cluster, and pre- or post-BCG treatment status. **D)** Histograms of entropy score per cell in the RNA (left) and ATAC (right) datasets. Shannon entropy is used as a measure of sample mixing in the local neighborhood (30 nearest neighbors) of a cell, calculated from the distribution of cells from each sample in that neighborhood. **E**) Gene ontology terms shared by HSPCs and cDCs based on significant differentially expressed genes in the combined RNA-seq dataset. **F**) Heat map of enrichment scores from ChromVAR for IRF transcription factors by cell type. **G)** Volcano plots of differential chromVAR motif accessibility in HSPCs, cDCs, and CD14+ monocytes. MeanDiff is the difference in average chromVAR score for accessible peaks post- *versus* pre- BCG, and p-values were determined by Wilcoxon rank-sum test. Significant motifs (p < 0.05) are colored according to relative enrichment up or down. H) Gene tracks depicting pseudobulk ATAC-seq accessibility peaks in HSPCs pre- (top, blue) and post-BCG (bottom, red) treatment for the HLA-C and HLA-DRB5 genes.

**Figure S2:**
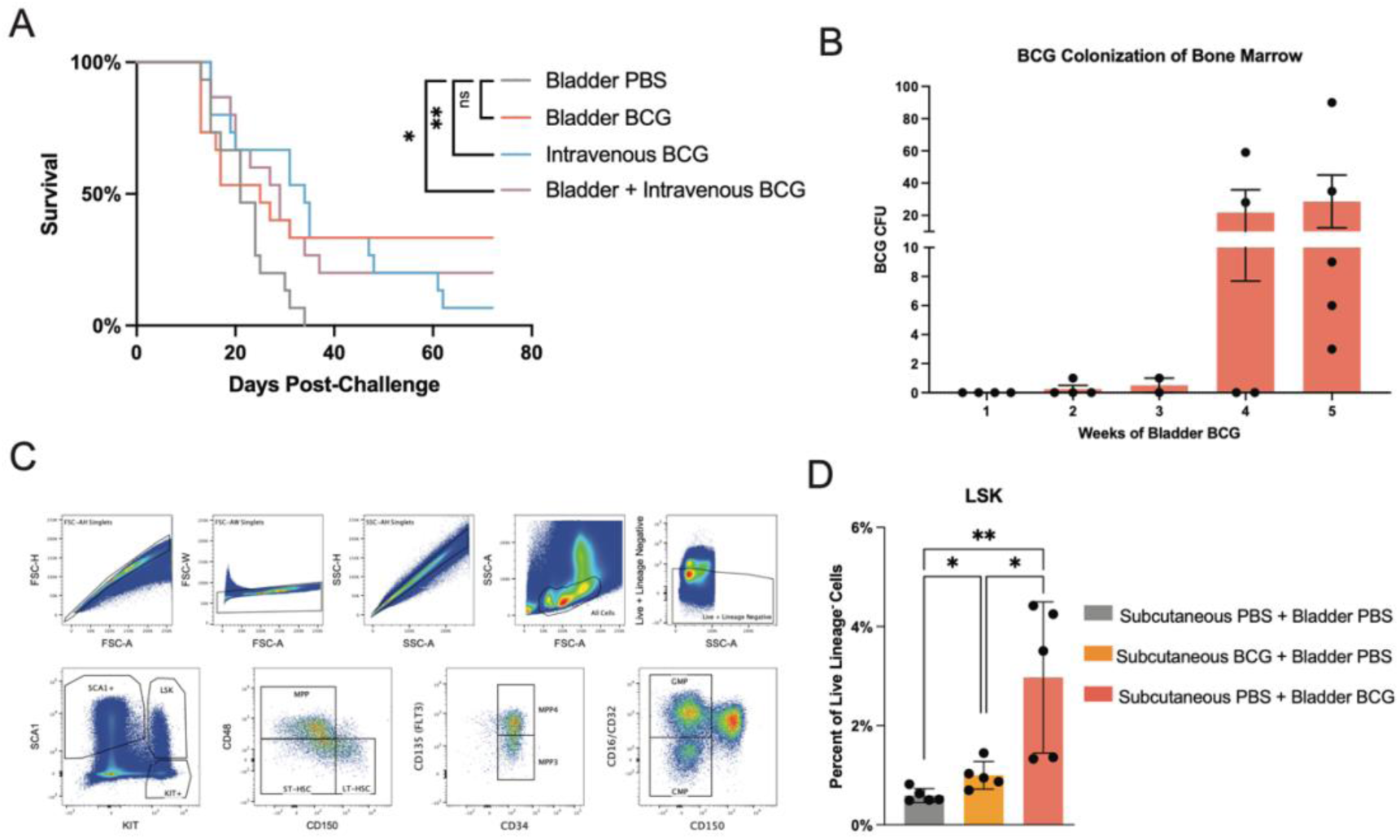
BCG colonizes the bone marrow after direct bladder administration and consequently improves survival against bladder tumors. A) Survival curve from the experiment depicted in Figure 2A. B) Individual bacterial load (CFU) data points from the time-course experiment depicted in Figure 2C. Error bars represent SEM. C) Gating hierarchy used to assess bone marrow hematopoietic stem and progenitor cell subsets in Figure 2. D) Quantification of Lineage- SCA1+ KIT+ (LSK) HSPC expansion following PBS, subcutaneous BCG, or bladder BCG administration. Error bars represent SD. P values for bar graphs were derived by Student’s t-test. P values for survival curves were derived by log-rank test. P > 0.05 = ns, P ≤ 0.05 = *, P ≤ 0.01 = **, P ≤ 0.001 = ***, P ≤ 0.0001 = ****.

**Figure S3:**
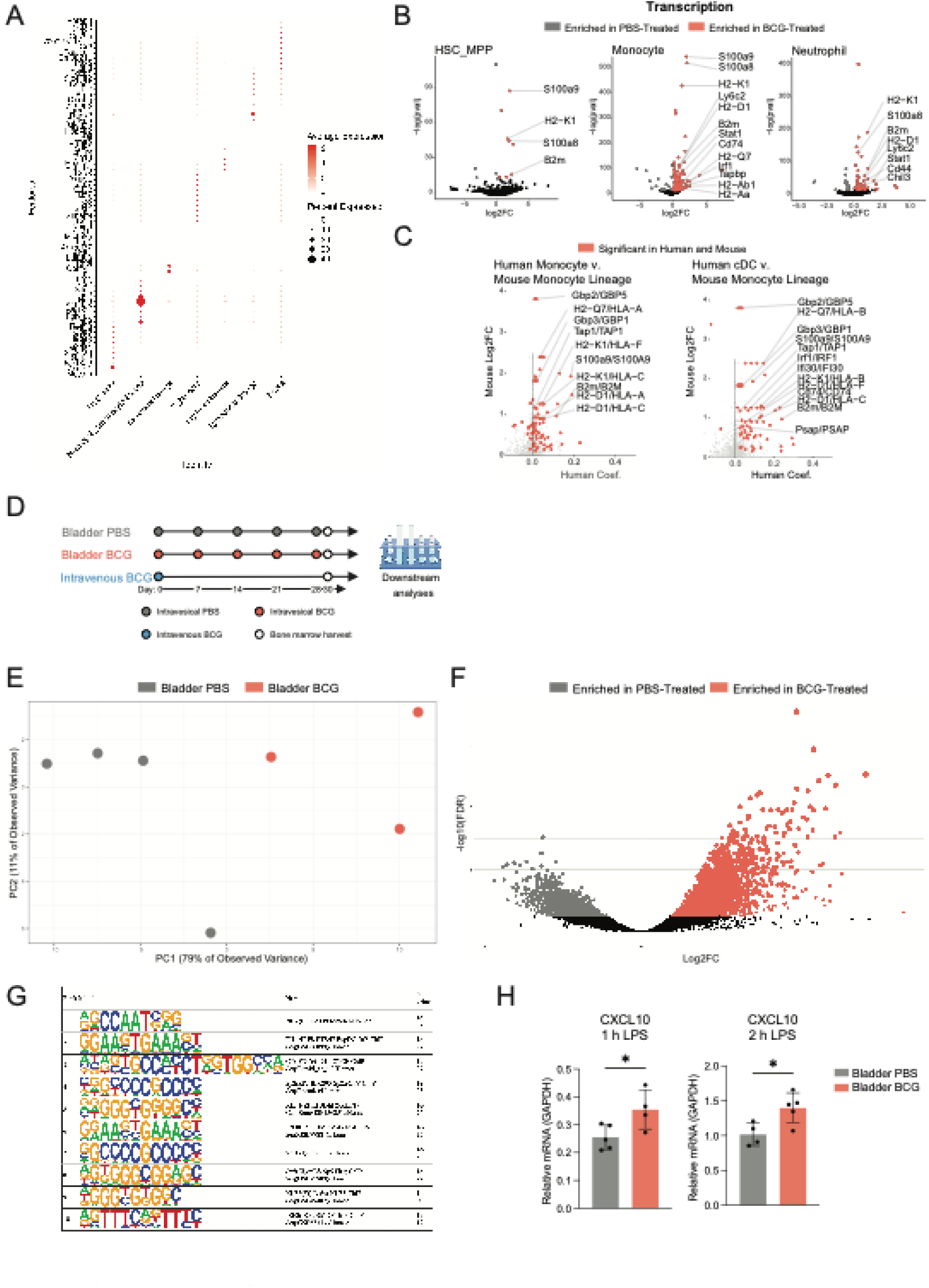
Bladder BCG administration alters chromatin accessibility in murine bone marrow LSK cells reflecting phenotypes in human BCG-treated bladder cancer patients. A) Dot plot of marker gene expression (columns) in each RNA PhenoGraph cluster (rows). Clusters are labeled by corresponding annotated cell type association, dots are colored by mean expression per cluster, and dot size corresponds to the fraction of cells in the cluster with non-zero expression of that gene. B) Volcano plots from snRNA-seq showing significant differentially expressed genes for HSC_MPP, monocytes, and neutrophils. C) Volcano plots depicting differentially expressed genes conserved between human and mouse cell subsets. Human monocyte *versus* mouse monocyte populations are shown on the left, human conventional dendritic cell *versus* mouse monocyte populations are shown on the right. Significant differentially expressed genes common to both humans and mice are highlighted in red. D) Experimental schematic. Mice were administered one of the following regimens beginning on Day 0: 5 weekly doses of bladder PBS, 5 weekly doses of bladder BCG, or a single dose of intravenous BCG. On Day 30, bone marrow was harvested and ATAC-seq was performed on sorted LSK cells. E) Principal Component Analysis (PCA) of ATAC-seq data from sorted LSKs. F) Volcano plot showing fold change of ATAC-seq peaks in LSKs from bladder PBS- *versus* bladder BCG-treated samples. G) HOMER motif enrichment analysis of differential ATAC-seq peaks in bladder PBS- *versus* bladder BCG-treated mice. H) LSK cells were sorted from PBS- or BCG-treated animals and differentiated into BMDMs using L929-conditioned medium. After 10 days, BMDMs were treated with LPS for 1 or 2 hours and CXCL10 mRNA was measured by RT-qPCR. P values for bar graphs were derived by Student’s t-test. P > 0.05 = ns, P ≤ 0.05 = *, P ≤ 0.01 = **, P ≤ 0.001 = ***, P ≤ 0.0001 = ****.

**Figure S4:**
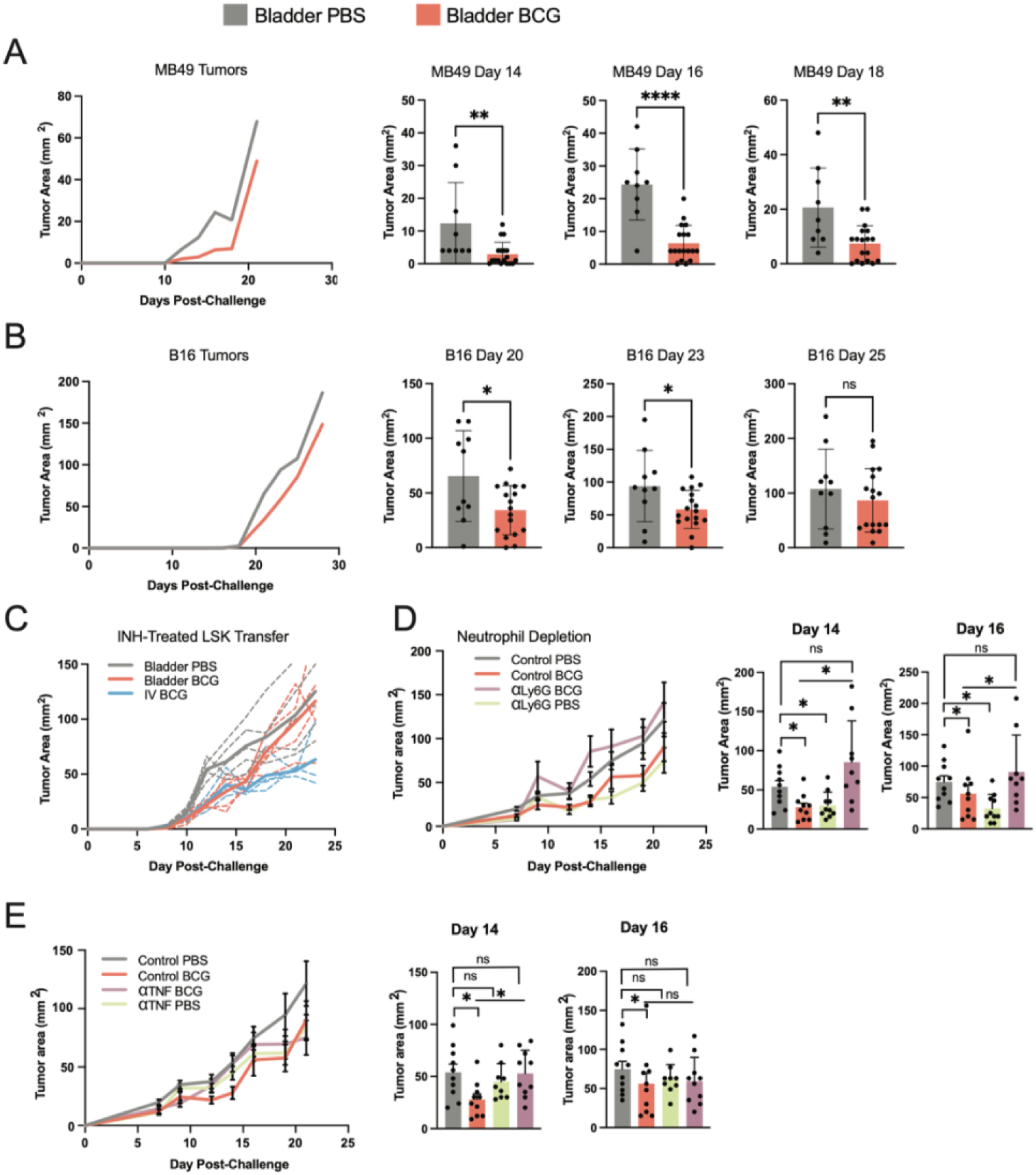
BCG reprogrammed HSPCs encode antitumor immunity to diverse tumor types. A) Tumor growth in naive irradiated recipient mice that received bone marrow from bladder PBS- or bladder BCG-treated donors, followed by subcutaneous challenge with MB49 tumors (left). Mid-curve time points are quantified for each cell line (right). B) Tumor growth in naive irradiated recipient mice that received bone marrow from bladder PBS- or bladder BCG-treated donors, followed by subcutaneous challenge with B16 melanoma tumors (left). Mid-curve time points are quantified (right). C) Subcutaneous tumor growth in naive irradiated recipient mice that received a transfer of sorted LSK cells expanded for 3 weeks on PVA media and treated with isoniazid for the entire duration. D) Subcutaneous tumor growth in naive irradiated recipient mice that received a transfer of bone marrow from bladder BCG donors (Red, Purple) or bladder PBS controls (Grey, Green). Animals were then treated with a Ly6G depleting antibody (Green, Purple) or an isotype control (Grey, Red). Mid-curve time points are quantified (right). E) Subcutaneous tumor growth in naive irradiated recipient mice that received a transfer of bone marrow from bladder BCG donors (Red, Purple) or bladder PBS controls (Grey, Green). Animals were then treated with a TNF-alpha blocking antibody (Green, Purple) or an isotype control (Grey, Red). Mid-curve time points are quantified (right). P values for bar graphs were derived by Student’s t-test. P > 0.05 = ns, P ≤ 0.05 = *, P ≤ 0.01 = **, P ≤ 0.001 = ***, P ≤ 0.0001 = ****.

**Figure S5:**
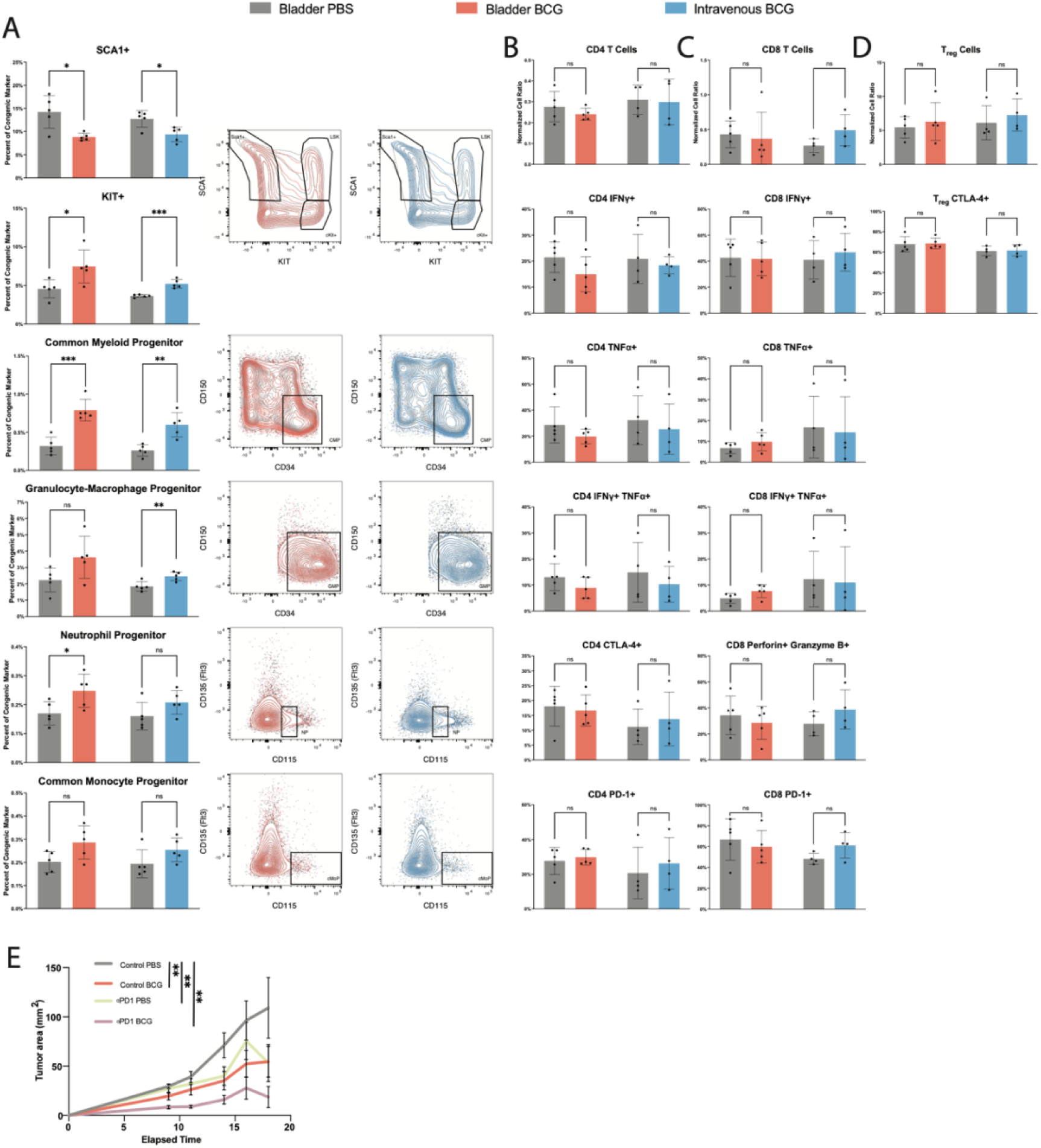
Comparisons of bone marrow HSPC subsets and tumor-infiltrating T cell subsets in BCG-experienced *versus* naive mixed bone marrow chimeric mice. A) Additional HSPC comparisons from the mixed bone marrow chimera experiment depicted in Figure 5A. Quantification of cell subsets from bladder or intravenous BCG-experienced *versus* naive origins are shown for Group 1 and Group 2, respectively (left), alongside representative flow plots (right). B-D) Quantification of T cell subsets from the mixed bone marrow chimera experiment depicted in Figure 5A. The following parameters were assessed for CD4 (B), CD8 (C), and Treg (D) subsets: abundance (fold change of cell frequency in the tumor compared to cell frequency in the spleen within each congenically-marked cell type), exhaustion (PD-1^+^ for CD4 and CD8 T cells, CTLA-4^+^ for CD4 T and Treg cells), and effector function (IFN-γ^+^, TNFɑ^+^, or IFN-γ^+^TNFɑ^+^ for CD4 and CD8 T cells). E) Subcutaneous tumor growth in naive irradiated recipient mice that received a transfer of bone marrow from bladder BCG donors (Red, Purple) or bladder PBS controls (Grey, Green). Animals were then treated with a PD1 blocking antibody (Green, Purple) or an isotype control (Grey, Red). P values for bar graphs were derived by Student’s t-test. P > 0.05 = ns, P ≤ 0.05 = *, P ≤ 0.01 = **, P ≤ 0.001 = ***, P ≤ 0.0001 = ****.

